# Chromatin alterations in the aging lung change progenitor cell activity

**DOI:** 10.1101/2021.07.15.452072

**Authors:** Samuel P. Rowbotham, Patrizia Pessina, Carolina Garcia de Alba Rivas, Jingyun Li, Irene G. Wong, Joon Yoon, Caroline Fahey, Aaron Moye, Joann Chongsaritsinsuk, Roderick Bronson, Shannan J. Ho Sui, Carla F. Kim

## Abstract

The lung contains multiple progenitor cell types that respond to damage, but how their responses are choreographed and why they decline with age is poorly understood. We report that histone H3 lysine 9 di-methylation (K9me2), mediated by histone methyltransferase G9a, regulates the dynamics of lung epithelial progenitor cells, and this regulation deteriorates with age. In aged mouse lungs, K9me2 loss coincided with lower frequency and activity of alveolar type 2 (AT2) cell progenitors. In contrast, K9me2 loss resulted in increased frequency and activity of multipotent progenitor cells with bronchiolar and alveolar potential (BASCs) and bronchiolar progenitors. K9me2 depletion in young mice through deletion or inhibition of G9a decreased AT2 progenitor activity and impaired alveolar injury regeneration. Conversely, K9me2 depletion increased chromatin accessibility of bronchiolar cell genes, increased BASC frequency and accelerated bronchiolar repair. K9me2 depletion also resulted in increased bronchiolar cell expression of the SARS-CoV2 receptor Ace2 in aged lungs. These data point to K9me2 and G9a as a critical regulator of the balance of lung progenitor cell regenerative responses and prevention of susceptibility to age-related lung diseases. These findings indicate that epigenetic regulation coordinates progenitor cell populations to expedite regeneration in the most efficient manner and disruption of this regulation presents significant challenges to lung health.

## Introduction

The mammalian lung is a large, complex organ, and as the only non-barrier system continuously exposed to the outside environment, the lung must protect itself and recover from external insults. Failure to do so has severe consequences; individual respiratory diseases account for five of the most common causes of death worldwide leading to 4 million annual deaths^1^ whilst 1 billion people live with acute or chronic respiratory conditions^2^. As lung disease imposes such an immense worldwide health burden, much effort has been expended in attempts to understand how lung epithelial progenitors maintain and regenerate functioning lung tissue. Lung function and regenerative capability declines during aging^3,4^, and aged lungs are the most vulnerable to respiratory diseases such as the chronic conditions COPD (Chronic Obstructive Pulmonary Disorder) and IPF (Idiopathic Pulmonary Fibrosis), and the novel SARS-CoV2 virus. Whilst observations suggest that regenerative capacity progressively decreases with age^5,6^, this has not been experimentally tested for lung progenitors that control regenerative responses. Further, it is not known how specific progenitor populations are affected, and how this translates to the specific disease vulnerabilities observed.

Epithelial progenitor cells are essential for long-term maintenance and renewal, as well as regeneration from acute injury. Multiple lung epithelial cell types have been described to function as progenitors, with functions largely associated with response to injury in the local environment of the trachea, airways or bronchioles, or the distal alveolar epithelium. Progenitor cell types defined include basal cells^7-9^, AT2 (Alveolar type 2) cells^10^, specialized AT2 cells^11,12^, club cells^13^, variant club cells^14,15^, BASCs (BronchioAlveolar Stem Cells)^16-18^, DASCs (Distal Alveolar Stem Cells)^19^, LNEPs (Lineage Negative Epithelial Progenitors)^20,21^ and neuroendocrine cells^22^. Some of these progenitors maintain well defined lineages, e.g. basal cells can give rise to club and ciliated cells. AT2 cells maintain the alveolar epithelium, sustaining the AT2 population and differentiating into AT1 cells^10,11^, bronchiolar cells including BASCs progenate club cells, and can also contribute to the alveolar epithelium, especially following acute alveolar injury^18,20,23^. It is not known why the lung has so many different progenitor cell types, and what controls their different behavior in homeostasis and injury response. To date, the progenitor functions of these cells have only been characterized in young animals, and it is also not known if and how these change with age, when regenerative function declines.

Numerous cellular processes are altered in the aging lung and have been implicated in degeneration, including proteostasis, senescence and mitochondrial function^3^. Considerably less is known about epigenetic changes, other than reports of increased transcriptional noise which generally implicate a loss of epigenetic fidelity^24^. Alongside the exhaustion of progenitor cells, the accumulation of epigenetic changes is also a hallmark of aging^25^, including the loss or disruption of repressive heterochromatin^26,27^. It has long been appreciated that epigenetic regulation is a key component controlling stem and progenitor cell function, with epigenetic modifications determining stem and differentiated cell fates^28^. Previous studies have identified roles for specific epigenetic regulators in lung function, lung diseases and lung progenitors, including the polycomb repressor Bmi1^29^, histone de-acetylases Hdac3^30^ and Sin3a^31^ and DNA methyltransferase Dnmt1^32^. It is clear from these studies that epigenetic regulators have essential roles in certain lung progenitors and at key developmental time points. However, it is not known if epigenetic regulation broadly controls how diverse lung progenitor populations are mobilized in response to different injuries.

One epigenetic regulator that has been implicated in various lung diseases is the histone methyltransferase G9a. G9a (also known as EHMT2) is responsible for catalyzing the di-methylation of histone H3 at lysine 9 (H3K9me2)^33^, a transcriptionally repressive modification^34^. G9a has been proposed to contribute to lung fibrosis by regulating fibroblasts^35^ and has been described as an oncogene in lung cancer^36,37^ as well as a regulator of lung cancer stem cells ^38^. Whilst lysine 9 methylation (K9me) is understood to generally decline in aging tissues and progerias^39,40^, very little is known about the dynamics of these histone modifications in the lung. As age is the most significant risk factor for chronic and acute lung diseases, this represents a substantial opportunity for inquiry.

Here we present evidence that G9a-mediated K9me2 can regulate lung cell fate and progenitor cell usage, which becomes dysregulated with age. In aged mouse lungs, global loss of K9me2 coincides with a reduction in the frequency of AT2 cells and increase in BASC frequency and bronchiolar progenitor activity. Depletion of K9me in young mice through genetics or inhibition of G9a reduces the activity of AT2 progenitors increases the number and activity of distal lung bronchiolar progenitors. This accelerates repair of bronchiolar injury but impairs regeneration of the alveoli, phenotypes seen in aging lungs. These effects are likely mediated through enhanced chromatin accessibility of bronchiolar secretory (club) cell genes. One consequence of this dysregulation is altered expression of the SARS-CoV-2 receptor Ace2, which is elevated in naturally K9me2-low aged club cells and K9me-depleted young club cells. These data point to K9me2 and G9a as a critical regulator of maintenance of the balance of lung progenitor cell regenerative responses and prevention of susceptibility to age-related lung diseases.

## Results

### Aged lung progenitor cells have altered frequency and activity

We used flow cytometry and histological analyses to deeply investigate lung progenitor cell abundance in the lung across different ages. On the gross morphological level, the lungs of young (2 months) and old (24 months) mice are indistinguishable (Figure S1a), but analysis of specific cell types revealed significant differences. Within the alveoli, AT2 cells have been demonstrated to maintain the epithelial compartment in young animals^10-12^. However, AT2 cells are significantly less frequent in the alveoli of old lungs than young (Figure 1a,b). Another progenitor population of the distal lung, the bronchiolalveolar stem cell (BASC) is capable of regenerating the alveoli and small airways^16-18^. In contrast with AT2 cells, BASCs, located at the bronchioalveolar duct junction, were significantly more frequent in old lungs than young (Figure 1c,d). Furthermore, the epithelial fractions which harbor these progenitors are also significantly altered in aged lungs, with the Epcam^+^ Sca-1^-^ population which consists of AT2 cells significantly smaller, and the Sca-1^+^ population, containing BASCs, significantly larger (Figure 1e,f).

**Figure 1.**
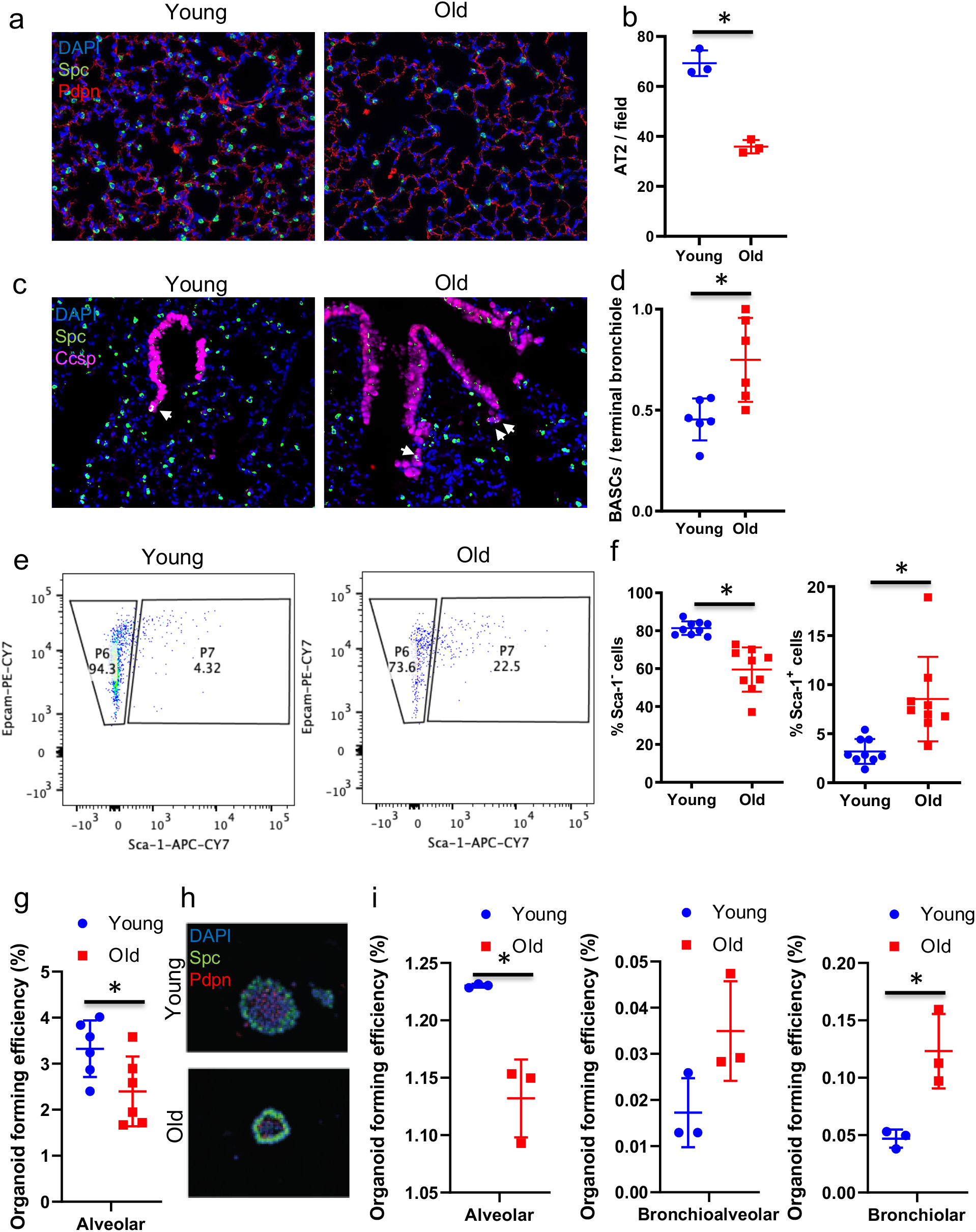
Lung progenitor populations alter in frequency and activity in aged mice. **(a)** Representative images of the alveolar space of young and old mice, immunostained for the indicated proteins. **(b)** Quantification of the number of AT2 cells per imaged alveolar field from young and old mice. ^*^=p<0.05, T-test. **(c)** Representative images of terminal bronchioles from young and old mice, immunostained for the indicated proteins. White arrows indicated BASCs. **(d)** Quantification of the number of BASCs per terminal bronchiole in young and old mice. ^*^=p<0.05, Mann-Whitney U-test. **(e)** Representative FACS plots from young and old mice, gated for single, live, Epcam^+^ CD45/^-^CD31^-^ cells. **(f)** Quantification of the proportion of Sca-1^-^ and Sca-1^+^ epithelial cells from young and old mice. ^*^=p<0.05, U-Test. **(g)** Quantification of the organoid forming efficiency of Sca-1^-^ AT2 cells isolated from young and old mice. ^*^=p<0.05, U-test **(h)** Representative images of organoids generated from Sca-1-cells from young and old mice, immunostained for the indicated proteins. **(i)** Quantification of the organoid forming efficiency of Sca-1+ cells from young and old mice. ^*^=p<0.05, U-test.

We next asked if the changes in cell numbers in old mice were also associated with altered progenitor cell activity by comparing young and old progenitor cells in our air-liquid interface organoid co-culture system^41^. In this system lung progenitors give rise to organoids resembling either the alveolar airspace (alveolar), the small airways (bronchiolar) and the bronchioalveolar duct junction (bronchioalveolar) (Figure S1b). Old Sca-1^-^ epithelial cells had a significantly lower organoid forming efficiency, giving rise to fewer alveolar organoids than the equivalent number of young cells (Figure 1g, S1c), and those that formed were smaller (Figure 1h). However, the proportion of organoids with cells positive for Pdpn, a marker of AT1 (Alveolar Type 1) cells, was not altered (Figure S1d), implying that the differentiation of AT2 into AT1 cells was not impaired. Old Sca-1^+^ epithelial cells, containing multipotent BASCs and other lung bronchiolar cells, did not show a significant difference in overall organoid forming efficiency (Figure S1e,f). However, old Sca-1+ progenitors generated significantly fewer alveolar organoids than young Sca-1+ cells (Figure 1i), whilst bronchioalveolar organoid forming efficiency was unchanged and bronchiolar organoid forming efficiency was significantly increased (Figure 1i). These results demonstrate that the pool of distal lung progenitors significantly changes with age, with fewer and less active alveolar progenitors and more frequent, and more active bronchiolar progenitors in old mice.

### Reduction of Histone H3 lysine 9 di-methylation in aged lungs is linked to reduced alveolar progenitor activity

We probed young and old lungs for gross changes in the amount of repressive chromatin modifications in epithelial cells, and we observed notable differences. Specifically, di-methylation of histone H3 lysine 9 was significantly reduced in both alveolar and bronchiolar cells in old compared to young lungs (Figure 2a,b). We have previously reported that depletion of H3K9me2, via inhibition of the histone methyltransferase G9a, strongly affects lung adenocarcinomas, altering their population dynamics to favor the expansion of aggressive, stem-like tumor propagating cells^38^. As there is evidence that tumors share regulatory mechanisms with healthy stem cells, we tested if depleting H3K9me2 by inhibiting G9a (hereafter G9ai) in young progenitors could alter their activity as we observed in their old counterparts (Figure 2c). We isolated young Sca-1^-^ and Sca-1^+^ progenitors. After 14 days of growth in organoid culture, H3K9me2 levels were significantly reduced in G9ai organoids compared to the vehicle control. (Figure S2a). Each of the major organoid types was observed with G9ai (Figure S2b) but with significantly different frequencies compared to the vehicle control (Figure S2c). Sca-1^-^ epithelial cells generated significantly fewer alveolar organoids with G9ai (Figure 2d), and those, as with organoids derived from old progenitors, were smaller with less branching (Figure S2b-c). The proportion of Pdpn^+^ organoids was also not altered (Figure S2b). Sca-1^+^ epithelial cells also generated significantly fewer alveolar organoids with G9ai, whilst bronchioalveolar organoid forming efficiency was unchanged and bronchiolar organoid forming efficiency was somewhat increased (Figure 2e).

**Figure 2.**
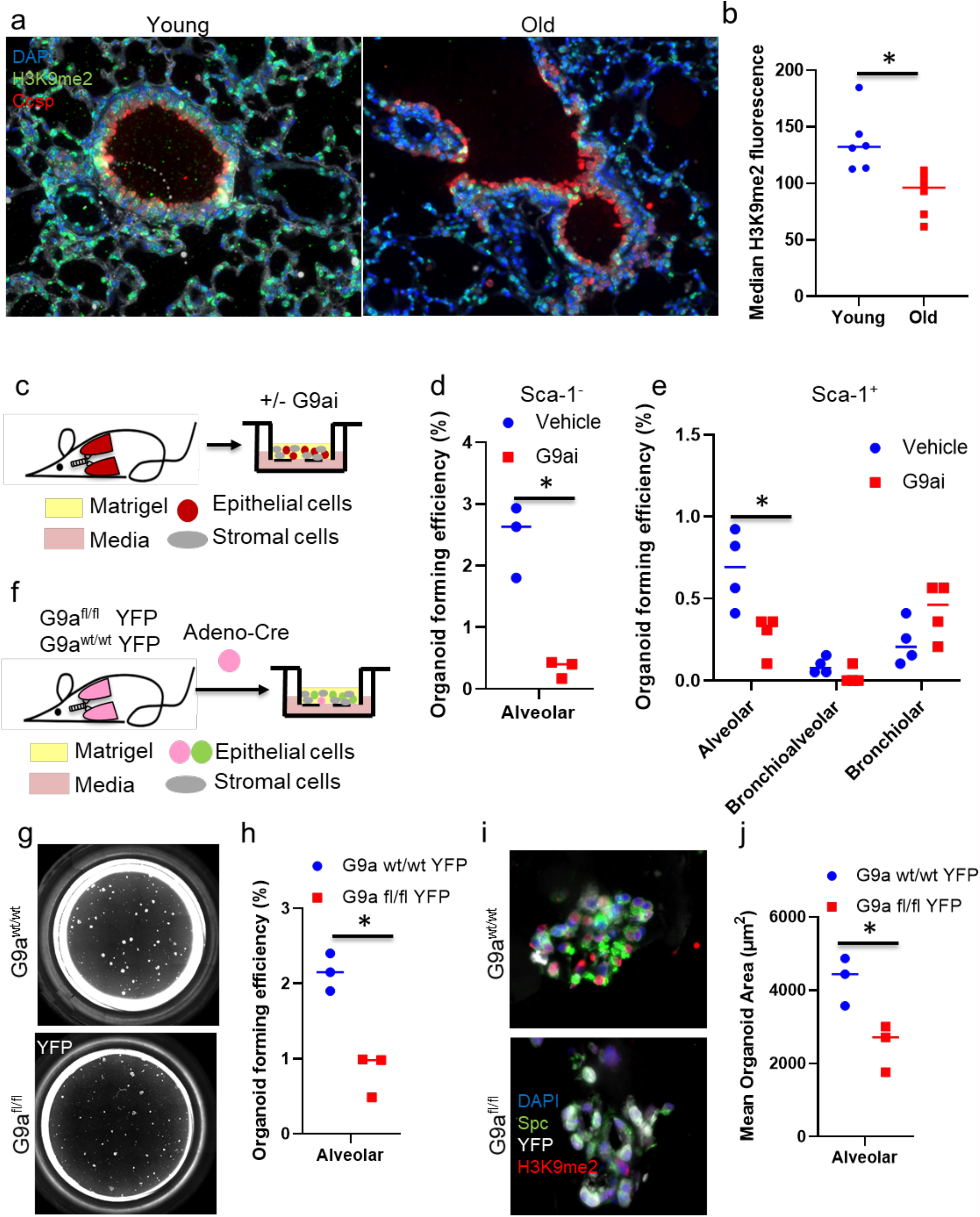
H3K9me2 levels are reduced in aged mice and H3K9me2 depletion in young mice impairs alveolar progenitor activity. **(a)** Representative images of young and old mouse lungs, immunostained for the indicated proteins. **(b)** Quantification H3K9me2 levels by fluorescent antibody imaging in young and old mouse lungs. ^*^=p<0.05, T-test. **(c)** Schematic of in vitro G9ai organoid experiment. **(d)** Quantification of organoid forming efficiencies from Sca-1^-^ progenitors +/- G9ai. ^*^=p<0.05, T-test. **(e)** Quantification of generation efficiencies of major organoid types from Sca-1^+^ progenitors +/- G9ai. ^*^=p<0.05, Sidak’s multiple comparison test. **(f)** Schematic of in vitro Adeno-Cre organoid experiment. **(g)** Representative images of YFP^+^ organoids derived from Adeno-Cre infected Sca-1^-^ progenitors from G9a^wt/wt^ and G9a^fl/fl^ mice, stained for YFP. **(h)** Quantification of organoid forming efficiencies from Adeno-Cre infected Sca-1^-^ progenitors from G9a^wt/wt^ and G9a^fl/fl^ mice. ^*^=p<0.05, T-test. **(i)** Representative images of organoids from Adeno-Cre infected Sca-1^-^ progenitors from G9a^wt/wt^ and G9a^fl/fl^ mice, immunostained for the indicated proteins. **(j)** Quantification of mean size of organoid derived from Adeno-Cre infected Sca-1^-^ progenitors from G9a^wt/wt^ and G9a^fl/fl^ mice. ^*^=p<0.05, T-test.

To determine if the effects on lung organoids were mediated through the activity of G9a in lung epithelial or stromal cells, we grew lung stromal cells +/- G9ai prior to epithelial cell co-culture (Figure S2e). Organoid forming efficiency for both Sca-1^-^ and Sca-1^+^ populations was unchanged between cultures with G9ai and control stromal cells (Figure S1f-g). Previously, we have reported that lung stromal cells support alveolar differentiation through a Thrombospondin1-BMP axis^41^. Relative expression of the stromal cell genes in this axis, Thbsp1, Bmp4, Hgf, as well as Tgfb which opposes alveolar differentiation, were not altered by G9ai (Figure S1h).

Finally, to verify that the phenotypes observed were due to H3K9me2 depletion and not off-target effects of the G9a inhibitor, we utilized a genetic model of G9a deletion. Sca-1^-^ progenitors were isolated from G9a^fl/fl lsl^YFP and ^lsl^YFP mice and incubated with Adeno-Cre virus to drive recombination of the floxed loci before plating in organoid culture (Figure 2f), as we have previously performed with ^lsl^KrasG12D mice^42^. After 14 days growth, we observed significantly fewer YFP^+^ alveolar organoids from the G9a^fl/fl^ progenitors (Figure 2g,h). Immunostaining of organoids confirmed that H3K9me2 was significantly depleted in YFP^+^ G9a^fl/fl^ organoids (Figure 2i, S2i). Furthermore, G9a^fl/fl^ organoids were also significantly smaller than WT organoids (Figure 2j). This confirms that depletion of H3K9me2 in young AT2 cells is sufficient to reduce their progenitor activity to that comparable of old progenitors, where repressive H3K9me2 modifications are significantly reduced.

### Alveolar injury repair is impaired in H3K9me2 depleted young mice

In mouse models of alveolar injury, old mice have severe difficulty in recovering from damage^43- 45^, which is similar to the chronic inability of the lungs to adequately regenerate in age-related diseases such as idiopathic pulmonary fibrosis (IPF). As H3K9me2-depleted young AT2 cells had reduced progenitor activity in vitro, like old AT2 cells, we tested if H3K9me2 depleted lungs would have similar problems with injury repair in vivo. We depleted H3K9me2 in vivo by bi-daily injections of 5 mg/kg of G9a inhibitor UNC0642^46^ and then subjected mice to bleomycin injury (Figure 3a). After two weeks, we observed significant reduction in H3K9me2 in the lung epithelia (Figure 3b). We analyzed the lungs at 21 days after injury, when repair of AT2 loss has begun, and 28 days when repair is more advanced. At 21 days there was moderate damage to both H3K9me2-depleted and control lungs with high variability between individual mice (Figure 3c,d). By day 28, much of the damage in control mice had been resolved, but damage persisted in G9ai mice (Figure 3d). Despite a slower resolution of lung damage, we surprisingly did not observe fewer AT2 cells in H3K9me2-depleted lungs (Figure S3a). However, when we measured AT2 cell proliferation by Ki67 we found there was significantly less AT2 cell proliferation in G9ai lungs (Figure 3e). In contrast to the reduction in proliferating AT2 cells, persistently damaged areas of H3K9me2-depleted lungs had significantly more BASC and club cells beyond the bronchioalveolar duct junction (Figure 3f,g). Intriguingly, these results suggest that loss of H3K9me2 may inhibit the normal repair of the alveoli by AT2 cells and force more of the regenerative burden onto the alterative progenitor populations of BASCs or club cells.

**Figure 3.**
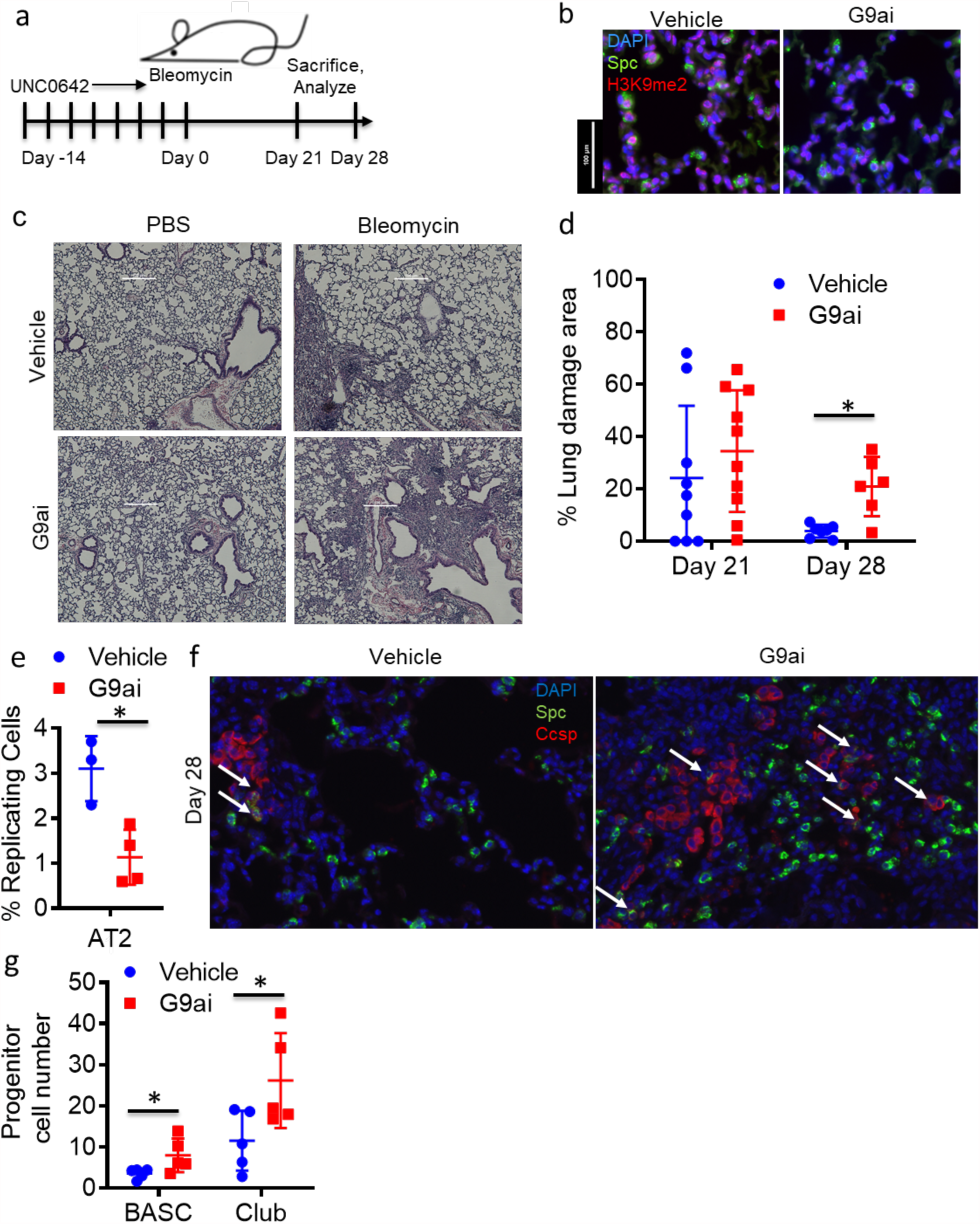
H3K9me2 depletion in young mice impairs alveolar injury repair and maintains aberrant bronchiolar cell expansion . **(a)** Schematic of G9ai and Bleomycin injury experiment. **(b)** Representative images of Representative images of alveolar airspaces of young mice following 14 days +/- G9ai, immunostained for the indicated proteins. **(c)** Representative H&E stained images of lungs from injured and injured G9ai and control mice. Scale bar=200 µM. **(d)** Quantification of the area of lung damage at the indicated timepoints following injury in control and G9ai mice. ^*^=p<0.05, t test. **(e)** Quantification of % of replicating AT2 cells in bleomycin damaged control and G9ai lungs. ^*^=p<0.05, t test **(f)** Representative images of damaged lung regions in G9ai and control mice, immunostained for the indicated proteins. Arrows demark Spc^+^ Ccsp^+^ cells. Scale bar = 100 µM. **(g)** Quantification of the number of BASC and extra-bronchiolar Ccsp+ cells per injured area in control and G9ai mice at day 28 following bleomycin injury. ^*^=p<0.05, U-Test.

### The bronchiolar progenitor population is expanded after in vivo H3K9me2 depletion

The presence of aberrantly expanded BASC and club cells in damaged lungs prompted us to study the populations of bronchiolar progenitors in H3K9me2 depleted lungs more closely in organoid culture (Figure 4a). As with alveolar cells, two weeks of in vivo G9ai significantly reduced levels of H3K9me2 in the bronchiolar epithelial cells (Figure 4b). As we observed with old lung epithelial, H3K9me2 reduction significantly increased the proportion of Sca-1^+^ cells (Figure 4c, S4a), although the corresponding decrease in Sca-1^-^ cells was too small to be considered statistically significant (Figure S4b). We found that the organoid forming efficiencies for bronchiolar and bronchioalveolar organoids appeared higher for the H3K9me2-depleted Sca-1^+^ progenitors, but the difference was not statistically significant (Figure S4c). The Sca-1^+^ population is heterogenous and we reasoned that with a more enriched population these differences may become clearer.

**Figure 4.**
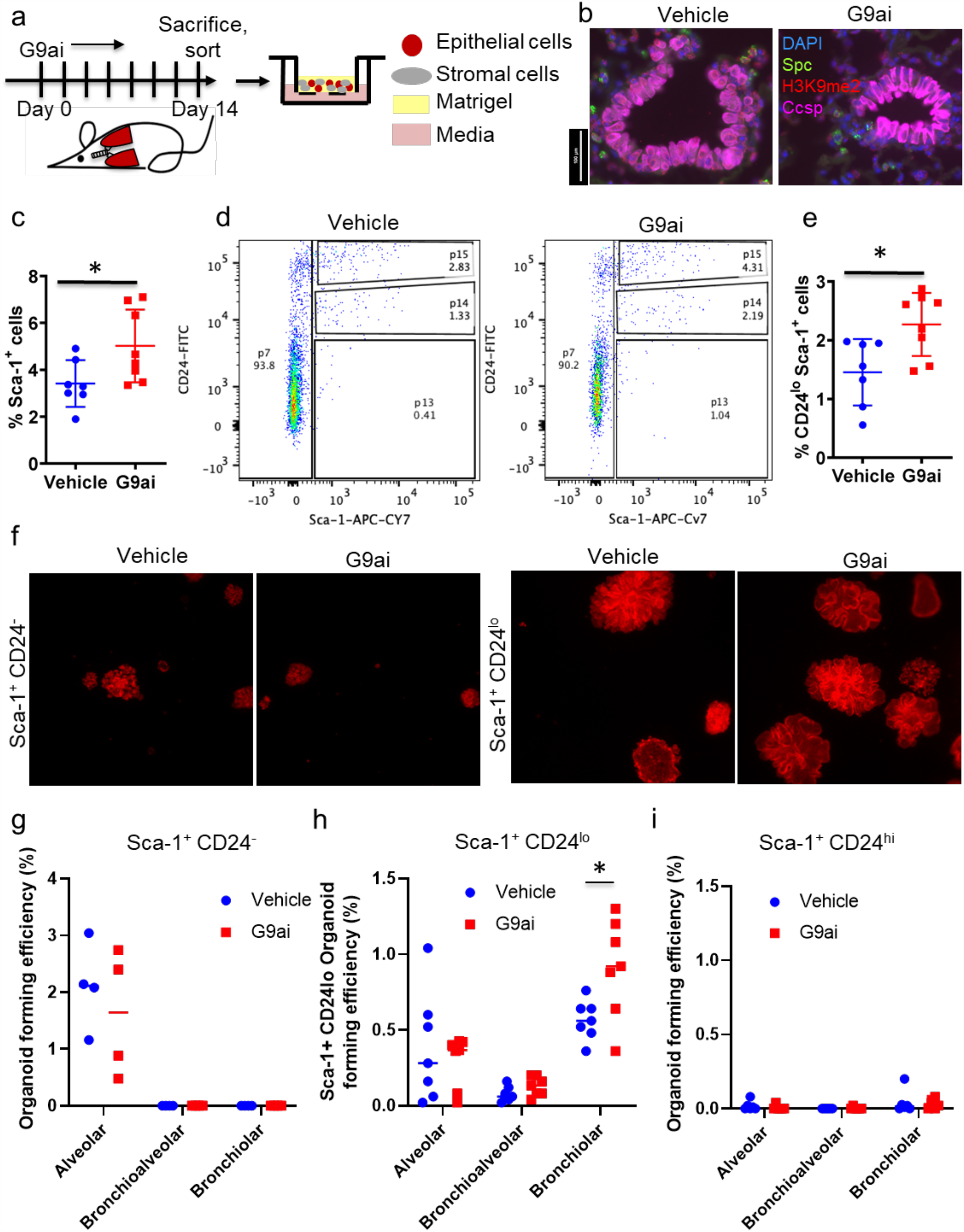
H3K9me2 depletion in young mice enriches for bronchiolar progenitors. **In vivo G9a inhibition enriches for bronchiolar progenitors**. **(a)** Schematic of in vivo G9ai organoid experiment. **(b)** Representative images of terminal bronchioles of young mice following 14 days +/- G9ai, immunostained for the indicated proteins. **(c)** Quantification of Sca-1^+^ fraction of lung epithelial cells, gated for single, live CD31/CD45^+^, Epcam^+^ cells. ^*^=p<0.05, T-test. **(d)** Representative FACs plots of G9ai and control isolated lungs, gated for single, live, CD31/CD45-, Epcam^+^ cells. **(e)** Quantification of the Sca-1^+^ CD24^lo^ fraction of lung epithelial cells, gated for single, live CD31/CD45^+^, Epcam^+^ cells. ^*^=p<0.05, T-test. **(f)** Representative fluorescent Images of day 14 organoid cultures from Sca-1^+^ CD24^-^ and Sca-1^+^ CD24^lo^ progenitors from G9ai and control mice. **(g-i)** Generation efficiencies of major organoid types from the **(g)** Sca-1^+^ CD24^-^, **(h)** Sca-1^+^ CD24^low^ and **(i)** Sca-1^+^ CD24^hi^ epithelial cells from G9ai and control mice. ^*^=p<0.05, Sidak’s multiple comparison test.

We further separated epithelial cell populations by using CD24, similar to a sorting strategy we had previously used^29^. Sorting G9ai and control epithelial cells again with CD24 produced three distinct Sca-1^+^ populations: CD24^hi^, CD24^lo^ and CD24^-^ (Figure 4d). QPCR of these populations for the lung epithelial cell markers *Sftpc, Scgb1a1*, and *Foxj1* suggested that the CD24^-^ population consists of AT2 cells, the CD24^lo^ is enriched in bronchiolar secretory (Club) cells and BASCs, and the CD24^hi^ in ciliated cells (Figure S4d-f). No population had significant expression of Krt5 or P63, implying that basal cells were absent from these preparations (data not shown). Of these populations, only the CD24^lo^ was significantly enriched in the G9ai lung epithelial population expressing EpCAM (Figure 4e, S4g,h). In organoid culture, the Sca-1^+^ CD24^-^ population gave rise only to alveolar organoids with a relatively high efficiency, (Figure 4f,g); this result was consistent with the gene expression data showing this population is comprised of AT2 cells. In contrast, the Sca-1^+^ CD24^lo^ cells formed bronchiolar and bronchioalveolar organoids with relatively high efficiency and significantly more bronchiolar organoids were generated from H3K9me2-depleted progenitors than controls (Figure 4f,h). The Sca-1^+^ CD24^hi^ population, the largest of the three, very rarely gave rise to organoids, as expected from a population enriched in ciliated cells (Figure 4i). These results demonstrate that the bronchiolar and bronchioalveolar progenitors are almost exclusively within the Sca-1^+^ CD24^lo^ population. The higher bronchiolar organoid forming efficiency of these cells, in combination with their higher proportion of the lung epithelium suggest that bronchiolar progenitor cells are significantly expanded by a reduction in H3K9me2 in young mice. Overall, our organoid experiments implied that G9a-mediated H3K9me2 has important functions regulating lung progenitors, as both old and H3K9me2-depleted young had a larger population of bronchiolar progenitors with higher progenitor activity than control young mice.

### Bronchiolar injury repair is accelerated after H3K9me2 depletion in young mice

We next probed the function of these progenitors in injury repair by subjecting H3K9me2-depleted and control young mice to naphthalene injury which selectively ablates club cells (Figure 5a). G9ai mice had significantly reduced H3K9me2 in the terminal bronchiolar epithelia (Figure S5a). Three days after injury, when repair of the airways is initiated, H3K9me2-depleted mice had regenerated significantly more club cells than controls, whereas club cells numbers are unchanged in the absence of injury (Figure 5b,c). This difference persisted a week following injury, when club cell numbers were almost at pre-injury levels in H3K9me2-depleted mice whereas regeneration was still incomplete in control mice (Figure 5c). As club cell numbers were restored more quickly with G9ai, we considered the possibility that more variant club cells with lower expression of *Cyp2f2*, the gene responsible for the toxic metabolism of Naphthalene, survive with G9ai and therefore re-population appears more advanced. However, chromatin accessibility of the *Cyp2f2* locus measured by ATAC-seq appears higher in G9ai bronchiolar progenitors (Figure S5b), arguing against this interpretation.

**Figure 5.**
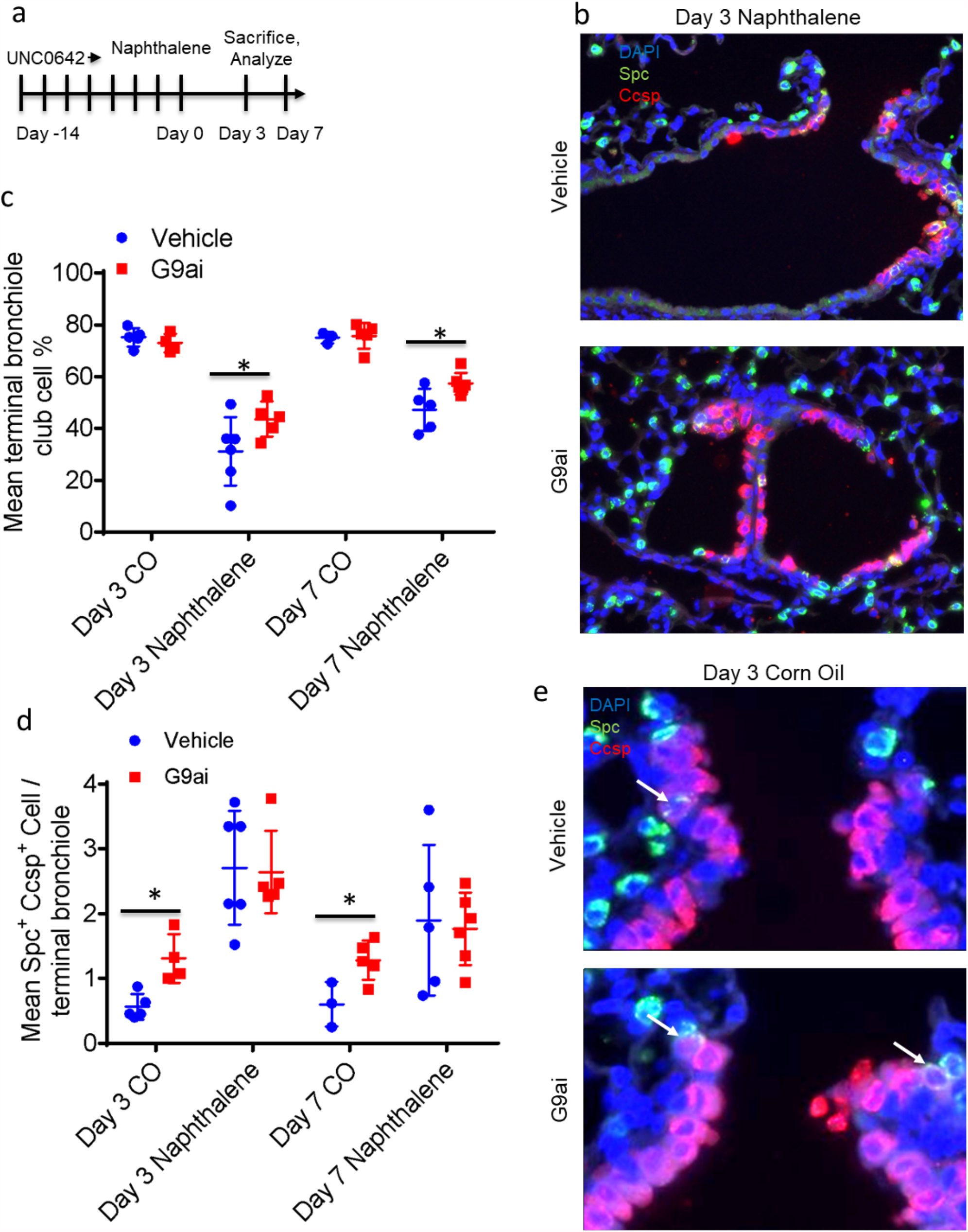
H3K9me2 depletion in young mice accelerates bronchiolar Injury repair. **(a)** Schematic of G9ai and Naphthalene injury experiment. **(b)** Representative images of damaged lung regions in G9ai and control mice, immunostained for the indicated proteins. Scale bar = 100 µM **(c)** Quantification of club cell loss and regeneration at the indicated timepoints in injured and uninjured control and G9ai mice. ^*^=p<0.05, Sidak’s multiple comparison test **(d)** Quantification of the number of BASCs per terminal bronchiole in injured and uninjured control and G9ai mice at the indicated timepoints. ^*^=p<0.05, Sidak’s multiple comparison test. **(e)** Representative images of terminal bronchioles in G9ai and control mice. Arrows demark Spc^+^ Ccsp^+^ BASCs. Scale bar = 100 µM.

As BASCs have been demonstrated to mediate a large proportion of repair of naphthalene-mediated damage at terminal bronchioles^16-18^, we wondered whether bronchiolar regeneration is accelerated by H3K9me2-depletion. We analyzed BASC numbers by quantifying Spc and Ccsp double positive cells at terminal bronchioles. We found that three days after injury, BASCs were expanded in both control and H3K9me2-depleted mice, with these numbers receding at day seven as repair progresses (Figure 5d). Interestingly, as we saw in old mice (Figure 1c,d), we found significantly more BASCs at terminal bronchioles in the uninjured G9ai mice vs controls (Figure 5d,e). Whilst BASCs were increased at G9ai terminal bronchioles, very few terminal bronchioles had BASCs adjacent to one another (Figure S5c). This suggests G9ai causes more cells to adopt the BASC fate, or resist differentiation into a club or AT2 cell, rather than the alternative explanation that G9ai causes existing BASCs to proliferate.

### H3K9me2 depletion increased the chromatin accessibility of young bronchiolar progenitors

In order to understand the mechanism of how G9a and H3K9me2 control the activity of lung epithelial progenitors, we performed ATAC-seq to determine the impact of active G9a on the accessibility of the chromatin landscape. ATAC-seq of isolated Sca-1^-^ (alveolar progenitors), Sca-1^+^ CD24^lo^ (bronchiolar progenitors) and Sca-1^+^ CD24^hi^ (non-progenitors) populations from control and H3K9me2-depleted mice revealed that accessible chromatin peaks were concentrated at the TSSs and 3’ UTRs of genes, as expected for mature epithelial cells (Figure S6a). As G9a-mediated H3K9me2 forms heterochromatin and represses transcription, we hypothesized the primary effect of G9ai H3K9me2-depletion would be to increase chromatin accessibility. Differences in peak signal enrichment between vehicle and G9ai were negligible in the Sca-1^-^ population, but signal enrichment was higher in G9ai Sca-1^+^ CD24^lo^ cells (Figure S6b). H3Kme2-depleted Sca-1^+^ CD24^lo^ cells had significantly more unique open chromatin peaks compared to vehicle (Figure 6a), but interestingly H3K9me2-depleted Sca-1^-^ cells had fewer (Figure S6c). Whilst unique peak intensity was almost identical in the Sca-1^-^ population, signal intensity was stronger in the H3K9me2-depleted Sca-1^+^ CD24^lo^ cells when compared to vehicle (Figure 6b). These analyses suggest that H3K9me2-depletion preferentially opens chromatin within bronchiolar progenitor populations.

**Figure 6.**
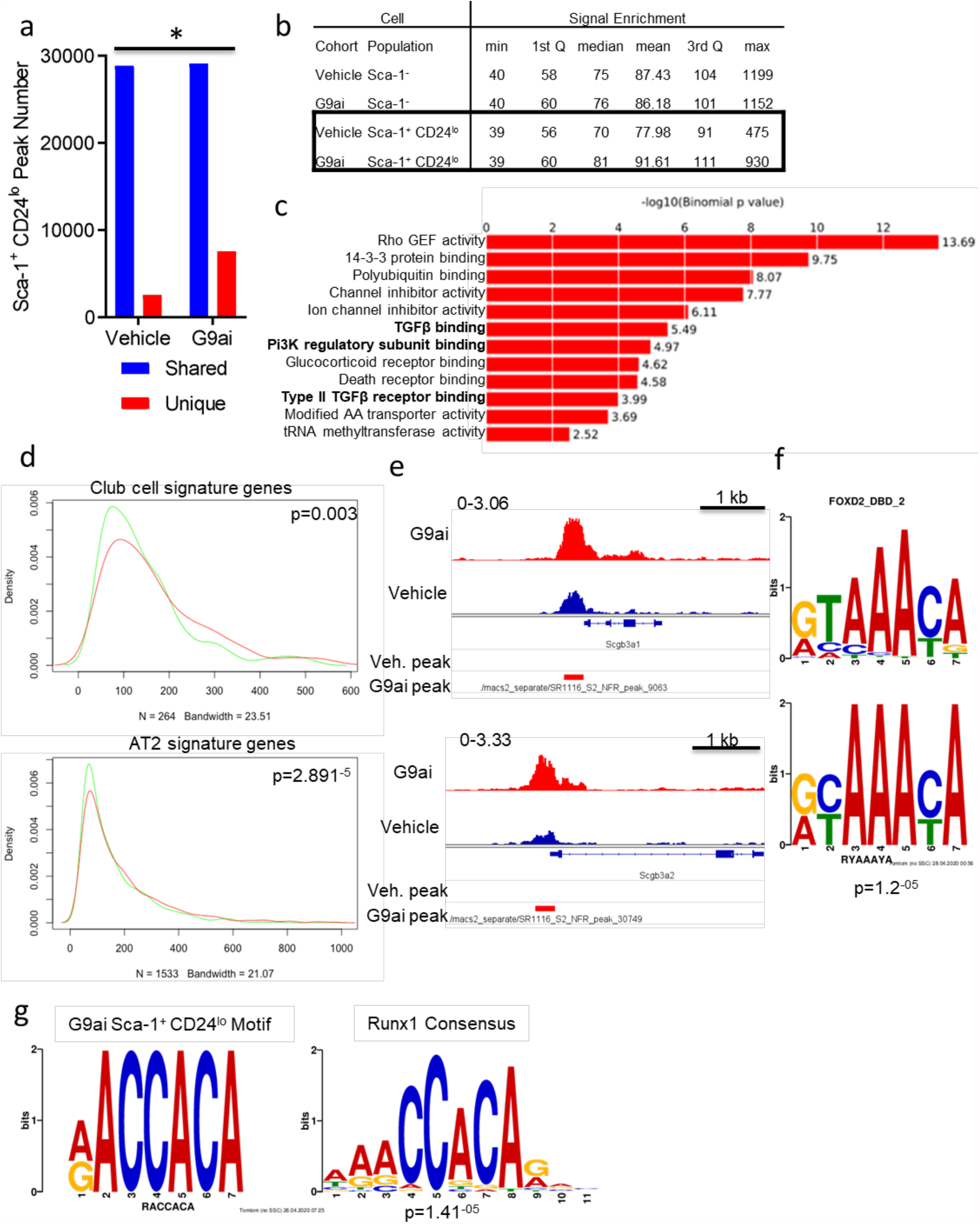
H3K9me2 depletion increases the chromatin accessibility of AT2 and Club cell genes in bronchiolar progenitors. **(a)** Quantification of common and unique ATAC-seq chromatin peaks in vehicle and G9ai Sca-1^+^ CD24^lo^ lung epithelial cells, p<0.05, χ^2^ test. **(b)** Table of chromatin peak signal enrichment of Sca-1^-^ and Sca-1^+^ CD24^lo^ cells from G9ai and vehicle lungs. **(c)** Table of significantly enriched GO Molecular Function terms associated with unique chromatin peaks from G9ai Sca-1^+^ CD24^lo^ cells. **(d)** Density plots of vehicle (green) and G9ai (red) open chromatin signal associated with differentially expressed genes of AT2 and club cell clusters from scRNA-seq of lung epithelial cells. P<0.005, Wilcoxon ranked-sum test. **(e)** Representative chromatin tracks of vehicle and G9ai Sca-1^+^ CD24^lo^ cells at highly specific club cell genes, with position of unique peaks from each cohort indicated. **(f)** Motif alignment between G9ai Sca-1^+^ CD24^lo^ unique peak enriched motif #1 and Foxd2 consensus. **(g)** Motif alignment between G9ai Sca-1^+^ CD24^lo^ unique peak enriched motif #7 and Runx1 consensus.

We next assessed the possible pathways that were influenced by altered chromatin accessibility in progenitor cell populations. We analyzed the genes associated with the chromatin regions that displayed more open accessibility from the Sca-1^-^ and Sca-1^+^ CD24^lo^ populations of H3K9me2-depleted and control mice, defined as peaks uniquely called in G9ai samples compared to vehicle using Gene Ontology analysis (Figure 6c, S6d). Amongst these GO terms were important cell signaling pathways, TGFB and Pi3K for H3K9me2-depleted Sca-1^+^ CD24^lo^ cells and MAP Kinase and Wnt for both H3K9me2-depleted and control Sca-1^-^ cells (Figure 6c, S6d,e). Differential expression of genes in these pathways could potentially mediate the differential progenitor activity of these cells.

To test for direct links between chromatin regulation and lung epithelial phenotypes, we analyzed the chromatin accessibility of genes that were highly expressed in AT2 cells or in club cells (bronchiolar secretory cells), using gene signatures we derived from single cell sequencing of Sca-1^-^ and Sca-1^+^ lung epithelial cells. We found significantly higher open chromatin signal enrichment amongst the club cell signature genes, and to a lesser extent the AT2 cell signature genes in the H3K9me2-depleted Sca-1^+^ CD24^lo^ cells (Figure 6d). Furthermore, 41/101 genes in the club cell signature were associated with a G9ai enriched peak versus 8/101 genes with a vehicle enriched peak, a significant enrichment (p<0.0001, χ^2^ test) (Table S1). This included the secretoglobins *Scgb1a1, Scgb3a2, Scgb3a1* which are amongst the most significantly upregulated club cell genes (Figure 6e). A smaller proportion of AT2 cell genes, 161/577 were associated with G9ai unique peaks and a comparable proportion, 54/577 with vehicle unique peaks (Table S2).

We also performed a motif enrichment analysis on each set of enriched peaks to determine if the binding sites of specific transcription factors (TFs) were represented in the differentially accessible chromatin upon H3K9me2 depletion. The highest enriched motif amongst the G9ai Sca-1^+^ CD24^lo^ peaks was highly similar to the consensus binding sequence of the Forkhead transcription factor Foxd2 (Figure 6f). Interestingly, a further highly enriched motif corresponded to Runx1, a transcription factor with increased expression in club cells (Figure 6g, Table S1.).

Whereas the top enriched motif, RYAAAYA most closely resembled the Foxd2 motif, Fox family members have very similar consensus binding sequences. Another Fox TF family member, Foxq1 is significantly upregulated in club cells (Table S1.) There is also strong homology between this motif and the core Foxq1 consensus RTAAACA (Figure S6f), raising the possibility that G9a-H3k9me2 could also govern the chromatin accessibility of this club cell TF to regulate bronchiolar cell gene expression.

### G9a inhibition recapitulated H3K9me2 reduction and changes in SARS-CoV2 receptor expression during aging

Our murine experiments suggested that the regenerative activity of lung progenitors is controlled by G9a-mediated H3K9me2 and disruption of this control leads to a bias towards bronchiolar progenitor activity and altered regenerative dynamics. Aged lungs are also more vulnerable to respiratory diseases, which has sharply been brought to international attention by the 2020 SARs-CoV2 pandemic. Data from recent studies suggest that the surface proteins required for SARs-CoV2 entry are concentrated in bronchiolar cells^47^ and immunostaining of the SARs-CoV2 receptor Ace2 in murine lungs confirmed this, with robust staining detected in club cells but not AT1, AT2 or ciliated cells (Figure 7a). We observed that expression was substantially higher in older mice, with a greater proportion of bronchiolar cells displaying luminal Ace2 staining (Figure 7b,c, S7a-c). When we analyzed the chromatin accessibility of the *Ace2* locus in control and H3K9me2-depleted young mice in the Sca-1^+^ CD24^lo^ bronchiolar progenitor population, we found increased accessibility in the H3K9me2-depleted cells (Figure 7d). Transcriptionally, *Ace2* expression was most pronounced in bronchiolar club cells (Figure S7b,c). Immunostaining of H3K9me2-depleted and control young mice revealed that like aged mice, the H3K9me2-depleted bronchioles had a significantly higher proportion of Ace2^+^ cells than controls (Figure 7e,f). These data suggest that H3K9me2 depletion through G9ai can recapitulate some of the phenotypes associated with aging lung epithelia and that epigenetic changes could underpin some of the vulnerabilities of the aged lung.

**Figure 7.**
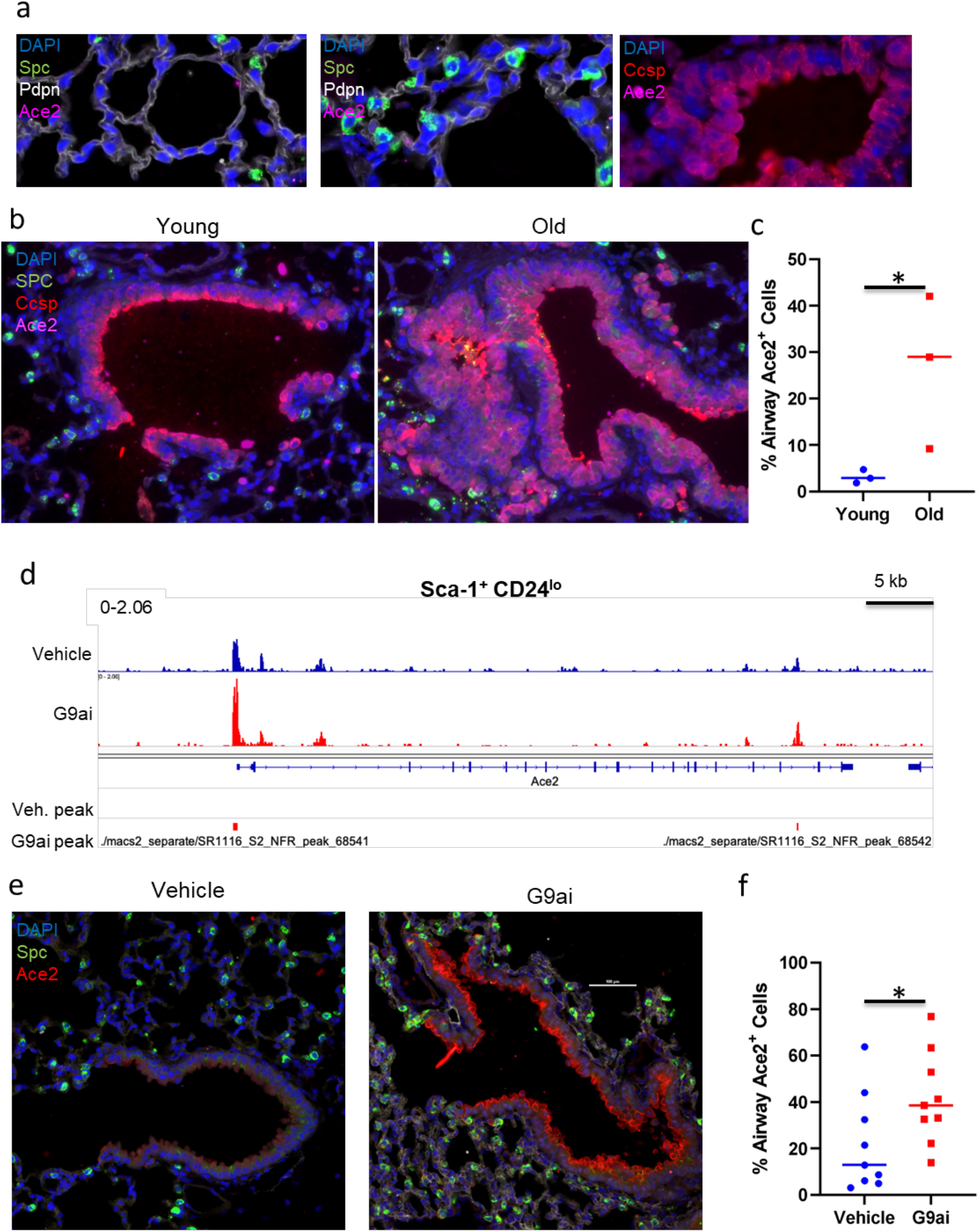
H3K9me2 depletion in young mice recapitulates aged SARS-CoV2 receptor expression. **(a)** Representative images of mouse alveolar airspaces and bronchioles, stained for the indicated proteins. **(b)** Representative images of bronchioles and surrounding alveoli in young and old mice, immunostained for the indicated proteins **(c)** Quantification of proportion of cells expressing Ace2 per imaged bronchiole of young and old mice. ^*^=p<0.05, T-test. **(d)** Representative chromatin tracks of vehicle and G9ai Sca-1^+^ CD24^lo^ cells at the Ace2 locus, with position of unique peaks from each cohort indicated. **(e)** Representative images of bronchioles and surrounding alveoli in control and G9ai mice, immunostained for the indicated proteins Scale bar = 100 µM. **(f)** Quantification of proportion of cells expressing Ace2 per imaged bronchiole of control and G9ai mice. ^*^=p<0.05, U test.

## Discussion

Our study provides insight into how rapid and accurate tissue repair is managed in an organ with multiple alternative progenitor cells, and how this changes with age. Most of our knowledge of lung regeneration has been derived from experiments in young adult mice, with some exceptions for the study of fibrotic disease^43-45^. Lineage tracing experiments in young mice have established that AT2 cells, or a specific progenitor subset of AT2 cells, preferentially replenish AT2 and AT1 cells in homeostasis and after acute alveolar damage^10-12^, whilst BASCs regenerate both the airways and alveoli around the duct junction following injury^17,18^. Our data suggest that changes in epigenetic modifications that occur with age contribute to the disruption of this process. We have shown that coincident with reduced global H3K9me2, aged mouse lungs have fewer AT2 cells and increased frequency of BASCs, the multipotent progenitors with alveolar and bronchiolar potential. AT2 cells from aged mice had reduced progenitor activity, as shown by decreased ability to form organoids. Further illustrating the altered balance of bronchiolar-alveolar homeostasis in aging, BASCs from old lungs produced fewer alveolar organoids and more bronchiolar organoids. Depletion of H3K9me2 in young mice through G9ai was sufficient to increase BASC frequency, bronchiolar progenitor activity, and the rate of regeneration of club cells at terminal bronchioles. Similarly, H3K9me2 depletion in young mice reduced AT2 progenitor activity, leading to impaired recovery from bleomycin-induced alveolar injury, which has been well documented in older mice^43-45^. H3K9me depletion increased the chromatin accessibility of genes expressed in bronchiolar cells, suggesting a mechanism for these effects.

These results therefore suggest a model of lung progenitor regulation whereby in young, healthy lungs, G9a-mediated H3K9me2 limits the chromatin accessibility of bronchiolar cell genes and restrains the ability of bronchiolar progenitor cells to respond to damage. In the event of injury to the alveoli, this layer of epigenetic transcriptional control would prevent excess activation of bronchiolar progenitors, perhaps limiting their regenerative contribution to the areas immediately surrounding the bronchioalveolar duct junction. This would allow the alveoli to be regenerated by AT2 progenitors, the preferred population to maintain AT2 and AT1 cells under homeostatic conditions. This response may reduce the likelihood of dysplastic airway-like cells persisting in the alveoli, a phenomenon known as bronchiolization, that is a feature of lung diseases such as IPF^48,49^. However, in aged lungs with the loss of K9me2-mediated transcriptional control, AT2 cells are less able to promptly respond to damage and hyperactivity of bronchiolar progenitors results in a slower repair process and a longer persistence of airway-like cells in the alveoli (Figure 8). It is interesting to consider if the increase in BASC numbers and progenitor activity are an adaptive response to aging, functioning as a reserve population to maintain and regenerate the alveoli that is increasingly relied upon as AT2 progenitor functions decrease. It is enticing that our findings could therefore explain the observations that cells with airway features are present in the alveolar space of IPF lungs.

**Figure 8.**
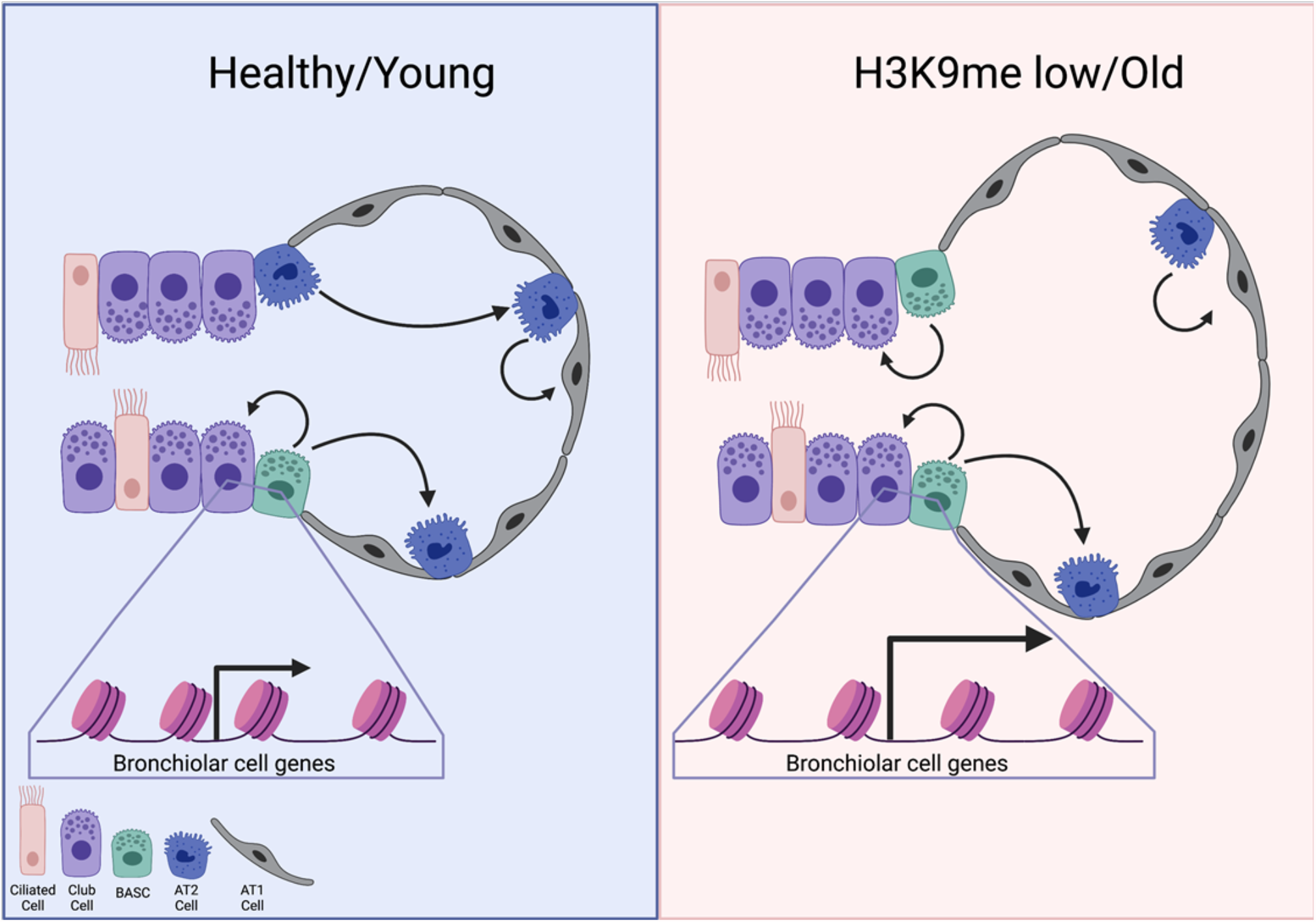
Chromatin changes in the aging lung change progenitor cell activity. In the young, healthy lung epithelium, G9a-mediated H3K9me2 restrains the transcriptional accessibility of bronchiolar cell genes, preventing excess activation of bronchiolar progenitors. This allows AT2 cells to efficiently regenerate the alveoli with the trade-off of slower replacement of club cells. In the aged, or artificially H3K9me depleted lung, expanded BASCs can regenerate the bronchioles more efficiently, but are less effective in compensating for the smaller number of active AT2 cells, leading to slower repair and the persistence of bronchiolar-like cells in the alveoli.

Whilst increased chromatin accessibility to airway cell genes presents a simple explanation for the bias towards airway regeneration with H3K9me2 loss, it is not clear if this is sufficient to also account for the decline in AT2 cell progenitor function. One possibility is that senescence (another hallmark of aging) increases in AT2 cells, although we did not detect any change in the chromatin accessibility of senescence genes in H3K9me2 depleted AT2 cells. The accumulation of senescent AT2 cells after bleomycin injury is predicted to drive pro-fibrotic signaling and limit repair and regeneration^50-52^. Interestingly, G9a has also been demonstrated have a pro-fibrosis effect in lung injury through its regulation of lung fibroblasts, preventing the return of myofibroblasts to a benign state^35^. In those experiments, G9ai after injury was insufficient to significantly aid recovery, possibly because of deleterious effects on AT2 cells, but specific deletion of G9a in fibroblasts improved fibrosis resolution. This contrasts with our results, where G9ai before injury impaired alveolar regeneration. A yet-to-be peer reviewed study suggests that G9a can regulate AT2 cells by a non-chromatin mediated mechanism^53^, although the significant changes we observe in H3K9me2 when G9a is inhibited or deleted does not lend weight to this particular interpretation.

The effects of aging on the lung are profound and multifaceted^3^, but the contribution of epigenetic changes to these phenotypes are chronically understudied. The effects of H3K9me loss in the aging process in other contexts are thought to have been primarily due to breakdown of pericentric chromatin that can lead to chromosomal instability and de-repression of repeat sequences. Our data suggest the possibility that de-regulation of progenitor activity and regeneration dynamics are also phenotypes of age-related H3K9me loss. Progenitor or stem cell exhaustion is often thought of as an independent facet or hallmark of aging, but our data suggest that this effect is downstream of more fundamental cellular processes, including epigenetic changes. Widespread loss of K9me heterochromatin in progeria has been shown to presage substantial losses in stem cell function^40^, leading to suggestions that this is a unified mechanism of aging. Our data lend weight to this interpretation that epigenetic changes are upstream of losses in stem cell function that appear with aging.

It is clear that progenitor activity and cell fate changes in aging lung have profound health consequences, and our work points to a particular vulnerability of aged lungs to respiratory infections, prominently highlighted by the 2020 SARS-CoV2 pandemic. Studies suggest that bronchiolar epithelial cells have significant expression of the receptors used by SARS-CoV2 to enter cells and are the likely source of initial transmission^47,54^. Our data suggest that in mice, Ace2 is one of several bronchiolar cell genes controlled by G9a, implying that loss of H3K9me with age will lead to increased Ace2 expression, which has been observed at the transcriptional level in patient samples^55^, and vulnerability to infection. Whilst Ace2 protein expression has been observed in CCSP^+^ cells in lung organoids^56^, to our knowledge this is the first clear demonstration that this protein is predominantly found in club cells in vivo. Studies using a mouse-modified SARS-CoV2 virus show strong infection of club cells in terminal bronchioles consistent with our findings^57^. The possibility of an epigenetic component to Covid-19 susceptibility is an intriguing prospect. Overall, our work supports further interrogation of G9a-mediated epigenetic regulation and other epigenetic mechanisms to utilize our understanding of lung progenitor activity for new interventions for the many respiratory diseases in which lung repair and regeneration are compromised.

## Methods

### Chemical Compounds

UNC0638 (Tocris) was reconstituted to 10 mM DMSO in stock and used at a final concentration of 250 nM in organoid & adherent cell culture. For cell culture experiments, cells were grown in the presence of the drug for 14 days with fresh media and drug supplied every 48 hrs. UNC0642 (Tocris) was reconstituted to 50 mg/ml in DMSO stock and diluted to 0.5 mg/ml in sterile filtered corn oil (Sigma) prior to use. Naphthalene (Sigma) was resuspended in sterile filtered corn oil at 27.5 mg/ml prior to use. Bleomycin (Sigma) was reconstituted at 20 mg/ml in sterile PBS.

### Mice & Tissues

Wild-type, Ds-red and G9a^fl/fl^ YFP mice were maintained in virus-free conditions on the C57/Black 6 background. G9a^fl/fl^ were kindly donated by Ann Schaefer from Mount Sinai School of Medicine. 18-month C57/Black 6 mice were acquired from Jackson laboratories and maintained in the BCH animal facility until 22-24 months of age for experimental use. All mouse experiments were approved by the AAALAC accredited BCH Animal Care and Use Committee and were performed in accordance with relevant institutional and national guidelines regulations. For In vivo G9ai experiments, mice were weighed and delivered 5 mg/kg UNC0642 in corn oil or vehicle control by Intraperitoneal injection, bi-daily at the same time of day for 14 days. Mice were monitored for weight loss and sickness. Lung tissue preparation was performed as previously described^41^ and sections were analyzed by at least two investigators including a pathologist. Lung injury from bleomycin was assessed by measuring the area of damaged tissue as determined by the pathologist and dividing by the total area of the lung.

### Flow Cytometry

Mice were euthanized with avertin overdose and lungs were dissected Tumors were dissected from the lungs of primary mice and tumor tissue was prepared as described^41^. Briefly, lungs were isolated, minced, digested rotating for 1h at 37°C with 2mg/ml Collagenase/Dispase (Roche) and then filtered twice (100 μm, then 40 μm) after a 5 minute incubation with 0.025 mg/mL DNase. Erythrocytes were lyzsed in a 90 s incubation, quenched with DMEM/FBS.

Single cell suspensions were stained using Sca1-APC-Cy7, CD45-APC, CD31-APC, Epcam-PE-Cy7, CD24-FITC with DAPI (Sigma) to visualize dead cells. All antibodies were incubated for 15 minutes at 1:100 dilutions. Cell sorting was performed with a FACS Aria II (Becton Dickson), and data were analyzed with FloJo software (Tree Star, Inc.).

### Stromal cell culture

Stromal cells were isolated largely as previously described^41^. Stromal cells were isolated from 2-to 4-week-old mice by negative selection with anti-CD45-conjugated magnetic beads and positive selection with anti-CD31-conjugated magnetic beads. CD31-positive cells were then amplified in a gelatin-coated culture plate for 3–5 days followed by reselection with anti-CD31-conjugated magnetic beads. For stromal cell G9ai, cells were cultured with 250 nm UNC0638 or vehicle control for 14 days, with media replenished every 2 days.

### Organoid Culture

Mouse organoid Culture was performed largely as previously described^41^, with 2500 epithelial cells plated per well. FACS-isolated lung epithelial cells were mixed with growth factor reduced Matrigel (Corning) containing lung stromal cells at 1×10^6^ cells/ml and plated in transwell plates. Cells were cultured for 14 days and analyzed. For in vitro G9ai, organoid cultures were grown with 250 nM UNC0638 or vehicle control for 14 days, with media replenished every 2 days. In vitro infection of lung epithelial cells was also performed as previously described^42^.

Ad5CMVCre or Ad5Empty adenoviruses (University of Iowa Virology Core) were diluted to 1×10^7^ PFU/ml in MTEC+ media. Freshly sorted lung epithelial cells were resuspended in virus solution at 1000 cells/m/ and incubated at for 1hr in 37 °C tissue culture incubated. Infected cells were washed 3x in PBS then plated with organoid cells in transwell plates.

### Immunofluorescence

Immunostaining of mouse lung tissue was performed as previously described (Rowbotham et al. 2018), using primary antibodies for pro-SPC (Abcam, 1:500), CCSP (T-18 and B-6, Santa Cruz, 1:200), acetylated α-tubulin (Sigma, 1:1000), H3K9me2 (ab1220, Abcam, 1:200 and Milipore, 1:100), Sox2 (149811-82, Invitrogen, 1:100), PDPN (Abcam, 1:200) Ace2, (PA5-48477, Invitrogen, 15 mg/ml). Secondary antibody staining included donkey α-rabbit 488, donkey α-goat 594, donkey α-goat 647, donkey α-mouse 594, goat α-mouse 647, goat α-rat 647, goat α-chicken 594 (1:200, Invitrogen). Sections were mounted and counterstained with DAPI. Immunostaining of organoids was also performed as previously described^41^. Matrigel plugs were mounted in histogel and sectioned, then immunostained as described for lung sections.

### RT-QPCR

QPCR was carried out as previously described^38^. Cultured cells or tissues were harvested and dissociated according to their specific protocols. RNA was isolated using either the Absolutely RNA Microprep Kit or Nanoprep Kit (Agilent), depending on cell numbers. cDNA was made using the SuperScript III kit (Invitrogen) and analyzed using TaqMan Assays (Applied Biosystems) with a StepOnePlus™ Real-Time PCR System (Applied Biosystems) and software as per the manufacturer’s recommendations. *Gapdh* (4352339E) was used as an endogenous control for normalization.

### Injury Experiments

For In vivo G9ai, mice were weighed and delivered 5 mg/kg UNC0642 in corn oil or vehicle control by intraperitoneal injection, bi-daily at the same time of day for 14 days. Naphthalene and bleomycin injury were performed as previously described^41^. In vivo lung injury experiments were conducted in 7- to 10-week-old mice that received naphthalene (275 mg/kg) via intraperitoneal injection or bleomycin (0.035 U/mice) via intratracheal injection. Mice were monitored for morbidity and were euthanized at the experimental timepoints indicated in the figures. Lungs were, inflated, fixed and processed for immunohistology.

### ATAC-seq

ATAC-seq of lung epithelial populations was performed using previously described methods^58^. Briefly, Sca-1^-^, Sca-1^+^ CD24^lo^ and Sca-1^+^ CD24^hi^ epithelial cells were isolated from the lungs of vehicle-treated or G9ai mice, and either 50,000 (Sca-1^-^) or 10,000 (Sca-1^+^ CD24^lo^ and Sca-1^+^ CD24^hi^) cells were used to for DNA Transposition and library generation. Quantified libraries were sequenced using an Illumina NextSeq 500 sequencer. Sequence quality was evaluated using FASTQC^59^. Reads were filtered and trimmed with Atropos^60^ if necessary. High quality reads were mapped to the mouse genome build (mm10) using Bowtie2^61^. After removing reads from mitochondrial DNA, we included properly paired reads with high mapping quality (MAPQ score >10)^62^ for further analysis. The ‘alignmentSieve’ function of Deeptools ^63^ and ‘sort’ and ‘index’ functions of Samtools^64^ were used to isolate fragments in the Nucleosome Free Regions (NFR) while considering the 9 bp shift (+4 in positive and -5 in negative strand) to account for Tn5 transposase binding as a dimer. Peaks were called using de-duplicated, uniquely mapped reads with MACS2^65^. Peaks with signal value >5 were retained. The final peak set quality was checked using the ChIPQC Bioconductor package^66^.

Differences in chromatin accessibility between groups were assessed using DiffBind ^67^. We did not observe significant batch effects due to sex or library preparation date during quality assessment. We used the default DESeq2^68^ incorporated in the DiffBind package for differential peak analysis. The peaks were filtered at the significance level of FDR < 0.05 and |log2 fold change| > 1. We additionally generated sets of “unique” and “shared” peak regions in comparisons of G9ai vs. vehicle using Bedtools^69^. The unique regions were annotated to nearby genes with ChIPseeker^70^ and analyzed for over-representation of Gene Ontology terms using GREAT^71^. Motif enrichment of unique regions was performed using MEME tools ^72^.

## Data Availability

All NGS sequencing data in this manuscript will be available from NCBI Geo. All other supporting data are available upon reasonable request of the corresponding author.

## Acknowledgements

We thank Ann Schaefer for providing the original G9a^fl/fl^ mice. We thank D. Kotton, for advice and critical reading of the manuscript and B. Stripp, M. Koenigshoff and members of the Kim lab for advice and discussions. We also thank R. Mathieu, M. Patinak and the Boston Children’s Hospital Flow Cytometry Core for assistance with flow cytometry experiments and The Harvard Medical School Biopolymers Facility for sequencing of NGS libraries.

## Author Contributions

S.P.R. designed, conceived, and performed the experiments and wrote the manuscript. P.P. and C.G.A. performed experiments with paired aged and young mice. J.L. and J.C. assisted with immunostaining experiments, I.G.W. and A.M. generated and analyzed scRNA-seq data. J.Y. and S.J.H.S. designed and performed the analysis of the ATAC-seq data. C.F. assisted with generation of G9a^fl/fl^ YFP mice. R.B. performed all histopathological analysis. C.F.K supervised the design and study and co-wrote the manuscript. All authors reviewed and edited the manuscript.

## References

1 Wang, H. et al. Global, regional, and national life expectancy, all-cause mortality, and cause-specific mortality for 249 causes of death, 1980–2015: a systematic analysis for the Global Burden of Disease Study 2015. The Lancet 388, 1459–1544, doi:https://doi.org/10.1016/S0140-6736(16)31012-1 (2016).

2 Mendis, S. et al. Global Status Report on noncommunicable diseases 2014. Geneva World Health Organization (2014).

3 Schneider, J. L. et al. The aging lung: Physiology, disease, and immunity. Cell 184, 1990–2019, doi:10.1016/j.cell.2021.03.005 (2021).

4 Navarro, S. & Driscoll, B. Regeneration of the Aging Lung: A Mini-Review. Gerontology 63, 270–280, doi:10.1159/000451081 (2017).

5 Schultz, M. B. & Sinclair, D. A. When stem cells grow old: phenotypes and mechanisms of stem cell aging. Development 143, 3–14, doi:10.1242/dev.130633 (2016).

6 Wells, J. M. & Watt, F. M. Diverse mechanisms for endogenous regeneration and repair in mammalian organs. Nature 557, 322–328, doi:10.1038/s41586-018-0073-7 (2018).

7 Rock, Jason R. et al. Notch-Dependent Differentiation of Adult Airway Basal Stem Cells. Cell Stem Cell 8, 639–648, doi:https://doi.org/10.1016/j.stem.2011.04.003 (2011).

8 Rock, J. R. et al. Basal cells as stem cells of the mouse trachea and human airway epithelium. Proceedings of the National Academy of Sciences 106, 12771–12775, doi:10.1073/pnas.0906850106 (2009).

9 Hong, K. U., Reynolds, S. D., Watkins, S., Fuchs, E. & Stripp, B. R. Basal cells are a multipotent progenitor capable of renewing the bronchial epithelium. Am J Pathol 164, 577–588, doi:10.1016/s0002-9440(10)63147-1 (2004).

10 Barkauskas, C. E. et al. Type 2 alveolar cells are stem cells in adult lung. J Clin Invest 123, 3025–3036, doi:10.1172/JCI68782 (2013).

11 Desai, T. J., Brownfield, D. G. & Krasnow, M. A. Alveolar progenitor and stem cells in lung development, renewal and cancer. Nature 507, 190–194, doi:10.1038/nature12930 (2014).

12 Zacharias, W. J. et al. Regeneration of the lung alveolus by an evolutionarily conserved epithelial progenitor. Nature 555, 251–255, doi:10.1038/nature25786 (2018).

13 Rawlins, E. L. et al. The role of Scgb1a1+ Clara cells in the long-term maintenance and repair of lung airway, but not alveolar, epithelium. Cell Stem Cell 4, 525–534, doi:10.1016/j.stem.2009.04.002 (2009).

14 Guha, A., Deshpande, A., Jain, A., Sebastiani, P. & Cardoso, W. V. Uroplakin 3a<sup>+</sup> Cells Are a Distinctive Population of Epithelial Progenitors that Contribute to Airway Maintenance and Post-injury Repair. Cell Reports 19, 246–254, doi:10.1016/j.celrep.2017.03.051 (2017).

15 Hong, K. U., Reynolds, S. D., Giangreco, A., Hurley, C. M. & Stripp, B. R. Clara cell secretory protein-expressing cells of the airway neuroepithelial body microenvironment include a label-retaining subset and are critical for epithelial renewal after progenitor cell depletion. Am J Respir Cell Mol Biol 24, 671–681, doi:10.1165/ajrcmb.24.6.4498 (2001).

16 Kim, C. F. B. et al. Identification of Bronchioalveolar Stem Cells in Normal Lung and Lung Cancer. Cell 121, 823–835 (2005).

17 Liu, Q. et al. Lung regeneration by multipotent stem cells residing at the bronchioalveolar-duct junction. Nature Genetics 51, 728–738, doi:10.1038/s41588-019-0346-6 (2019).

18 Salwig, I. et al. Bronchioalveolar stem cells are a main source for regeneration of distal lung epithelia in vivo. The EMBO Journal 38, e102099, doi:10.15252/embj.2019102099 (2019).

19 Zuo, W. et al. p63(+)Krt5(+) distal airway stem cells are essential for lung regeneration. Nature 517, 616–620, doi:10.1038/nature13903 (2015).

20 Kathiriya, J. J., Brumwell, A. N., Jackson, J. R., Tang, X. & Chapman, H. A. Distinct Airway Epithelial Stem Cells Hide among Club Cells but Mobilize to Promote Alveolar Regeneration. Cell Stem Cell 26, 346-358.e344, doi:10.1016/j.stem.2019.12.014 (2020).

21 Vaughan, A. E. et al. Lineage-negative progenitors mobilize to regenerate lung epithelium after major injury. Nature 517, 621–625, doi:10.1038/nature14112 (2015).

22 Ouadah, Y. et al. Rare Pulmonary Neuroendocrine Cells Are Stem Cells Regulated by Rb, p53, and Notch. Cell 179, 403-416.e423, doi:10.1016/j.cell.2019.09.010 (2019).

23 Tropea, K. A. et al. Bronchioalveolar stem cells increase after mesenchymal stromal cell treatment in a mouse model of bronchopulmonary dysplasia. Am J Physiol Lung Cell Mol Physiol 302, L829–837, doi:10.1152/ajplung.00347.2011 (2012).

24 Angelidis, I. et al. An atlas of the aging lung mapped by single cell transcriptomics and deep tissue proteomics. Nature Communications 10, 963, doi:10.1038/s41467-019-08831-9 (2019).

25 López-OtÍn, C., Blasco, M. A., Partridge, L., Serrano, M. & Kroemer, G. The Hallmarks of Aging. Cell 153, 1194–1217, doi:10.1016/j.cell.2013.05.039 (2013).

26 Nardini, C. et al. The epigenetics of inflammaging: The contribution of age-related heterochromatin loss and locus-specific remodelling and the modulation by environmental stimuli. Seminars in Immunology 40, 49–60, doi:https://doi.org/10.1016/j.smim.2018.10.009 (2018).

27 Tsurumi, A. & Li, W. X. Global heterochromatin loss: a unifying theory of aging? Epigenetics 7, 680–688, doi:10.4161/epi.20540 (2012).

28 Chen, T. & Dent, S. Y. Chromatin modifiers and remodellers: regulators of cellular differentiation. Nat Rev Genet 15, 93–106, doi:10.1038/nrg3607 (2014).

29 Zacharek, S. J. et al. Lung stem cell self-renewal relies on Bmi1-dependent control of expression at imprinted loci. Cell stem cell 9, 272–281, doi:10.1016/j.stem.2011.07.007 (2011).

30 Wang, Y. et al. HDAC3-Dependent Epigenetic Pathway Controls Lung Alveolar Epithelial Cell Remodeling and Spreading via miR-17-92 and TGF-β Signaling Regulation. Dev Cell 36, 303–315, doi:10.1016/j.devcel.2015.12.031 (2016).

31 Yao, C. et al. Sin3a regulates epithelial progenitor cell fate during lung development. Development 144, 2618–2628, doi:10.1242/dev.149708 (2017).

32 Liberti, D. C. et al. Dnmt1 is required for proximal-distal patterning of the lung endoderm and for restraining alveolar type 2 cell fate. Developmental Biology 454, 108–117, doi:https://doi.org/10.1016/j.ydbio.2019.06.019 (2019).

33 Tachibana, M. et al. G9a histone methyltransferase plays a dominant role in euchromatic histone H3 lysine 9 methylation and is essential for early embryogenesis. Genes & Development 16, 1779–1791, doi:10.1101/gad.989402 (2002).

34 Tachibana, M. et al. Histone methyltransferases G9a and GLP form heteromeric complexes and are both crucial for methylation of euchromatin at H3-K9. Genes Dev 19, 815–826, doi:10.1101/gad.1284005 (2005).

35 Ligresti, G. et al. CBX5/G9a/H3K9me-mediated gene repression is essential to fibroblast activation during lung fibrosis. JCI Insight 5, doi:10.1172/jci.insight.127111 (2019).

36 Chen, M.-W. et al. H3K9 Histone Methyltransferase G9a Promotes Lung Cancer Invasion and Metastasis by Silencing the Cell Adhesion Molecule Ep-CAM. Cancer Research 70, 7830–7840, doi:10.1158/0008-5472.can-10-0833 (2010).

37 Zhang, K. et al. Targeting histone methyltransferase G9a inhibits growth and Wnt signaling pathway by epigenetically regulating HP1α and APC2 gene expression in non-small cell lung cancer. Mol Cancer 17, 153, doi:10.1186/s12943-018-0896-8 (2018).

38 Rowbotham, S. P. et al. H3K9 methyltransferases and demethylases control lung tumor-propagating cells and lung cancer progression. Nature Communications 9, 4559, doi:10.1038/s41467-018-07077-1 (2018).

39 Djeghloul, D. et al. Age-Associated Decrease of the Histone Methyltransferase SUV39H1 in HSC Perturbs Heterochromatin and B Lymphoid Differentiation. Stem Cell Reports 6, 970–984, doi:https://doi.org/10.1016/j.stemcr.2016.05.007 (2016).

40 Zhang, W. et al. A Werner syndrome stem cell model unveils heterochromatin alterations as a driver of human aging. Science 348, 1160, doi:10.1126/science.aaa1356 (2015).

41 Lee, J.-H. et al. Lung stem cell differentiation in mice directed by endothelial cells via a BMP4-NFATc1-Thrombospondin-1 axis. Cell 156, 440–455, doi:10.1016/j.cell.2013.12.039 (2014).

42 Dost, A. F. M. et al. Organoids Model Transcriptional Hallmarks of Oncogenic KRAS Activation in Lung Epithelial Progenitor Cells. Cell Stem Cell 27, 663-678.e668, doi:10.1016/j.stem.2020.07.022 (2020).

43 Hecker, L. et al. Reversal of persistent fibrosis in aging by targeting Nox4-Nrf2 redox imbalance. Sci Transl Med 6, 231ra247, doi:10.1126/scitranslmed.3008182 (2014).

44 Huang, W. T. et al. Plasminogen activator inhibitor 1, fibroblast apoptosis resistance, and aging-related susceptibility to lung fibrosis. Experimental gerontology 61, 62–75, doi:10.1016/j.exger.2014.11.018 (2015).

45 Sueblinvong, V. et al. Predisposition for disrepair in the aged lung. The American journal of the medical sciences 344, 41–51, doi:10.1097/MAJ.0b013e318234c132 (2012).

46 Liu, F. et al. Discovery of an in Vivo Chemical Probe of the Lysine Methyltransferases G9a and GLP. Journal of Medicinal Chemistry 56, 8931–8942, doi:10.1021/jm401480r (2013).

47 Sungnak, W. et al. SARS-CoV-2 entry factors are highly expressed in nasal epithelial cells together with innate immune genes. Nature Medicine 26, 681–687, doi:10.1038/s41591-020-0868-6 (2020).

48 Plantier, L. et al. Ectopic respiratory epithelial cell differentiation in bronchiolised distal airspaces in idiopathic pulmonary fibrosis. Thorax 66, 651–657, doi:10.1136/thx.2010.151555 (2011).

49 Seibold, M. A. et al. The idiopathic pulmonary fibrosis honeycomb cyst contains a mucocilary pseudostratified epithelium. PLoS One 8, e58658–e58658, doi:10.1371/journal.pone.0058658 (2013).

50 Parimon, T., Yao, C., Stripp, B. R., Noble, P. W. & Chen, P. Alveolar Epithelial Type II Cells as Drivers of Lung Fibrosis in Idiopathic Pulmonary Fibrosis. Int J Mol Sci 21, 2269, doi:10.3390/ijms21072269 (2020).

51 Xu, Y. et al. Single-cell RNA sequencing identifies diverse roles of epithelial cells in idiopathic pulmonary fibrosis. JCI insight 1, e90558–e90558, doi:10.1172/jci.insight.90558 (2016).

52 Selman, M. & Pardo, A. The leading role of epithelial cells in the pathogenesis of idiopathic pulmonary fibrosis. Cell Signal 66, 109482, doi:10.1016/j.cellsig.2019.109482 (2020).

53 Pribluda, A. et al. G9a methyltransferase governs cell identity in the lung and is required for KRAS G12D tumor development and propagation. bioRxiv, 2020.2004.2020.050328, doi:10.1101/2020.04.20.050328 (2020).

54 Ziegler, C. G. K. et al. SARS-CoV-2 Receptor ACE2 Is an Interferon-Stimulated Gene in Human Airway Epithelial Cells and Is Detected in Specific Cell Subsets across Tissues. Cell 181, 1016-1035.e1019, doi:https://doi.org/10.1016/j.cell.2020.04.035 (2020).

55 Lukassen, S. et al. SARS-CoV-2 receptor ACE2 and TMPRSS2 are primarily expressed in bronchial transient secretory cells. Embo j 39, e105114, doi:10.15252/embj.20105114 (2020).

56 Salahudeen, A. A. et al. Progenitor identification and SARS-CoV-2 infection in human distal lung organoids. Nature 588, 670–675, doi:10.1038/s41586-020-3014-1 (2020).

57 Leist, S. R. et al. A Mouse-Adapted SARS-CoV-2 Induces Acute Lung Injury and Mortality in Standard Laboratory Mice. Cell 183, 1070-1085.e1012, doi:10.1016/j.cell.2020.09.050 (2020).

58 Corces, M. R. et al. An improved ATAC-seq protocol reduces background and enables interrogation of frozen tissues. Nature Methods 14, 959–962, doi:10.1038/nmeth.4396 (2017).

59 Andrews, S. FastQC: a quality control tool for high throughput sequence data. http://www.bioinformatics.babraham.ac.uk/projects/fastqc (2010).

60 Didion, J. P., Martin, M. & Collins, F. S. Atropos: specific, sensitive, and speedy trimming of sequencing reads. PeerJ 5, e3720, doi:10.7717/peerj.3720 (2017).

61 Langmead, B., Trapnell, C., Pop, M. & Salzberg, S. L. Ultrafast and memory-efficient alignment of short DNA sequences to the human genome. Genome biology 10, R25, doi:10.1186/gb-2009-10-3-r25 (2009).

62 Tarasov, A., Vilella, A. J., Cuppen, E., Nijman, I. J. & Prins, P. Sambamba: fast processing of NGS alignment formats. Bioinformatics 31, 2032–2034, doi:10.1093/bioinformatics/btv098 (2015).

63 RamÍrez, F., Dündar, F., Diehl, S., Grüning, B. A. & Manke, T. deepTools: a flexible platform for exploring deep-sequencing data. Nucleic acids research 42, W187–191, doi:10.1093/nar/gku365 (2014).

64 Li, H. et al. The Sequence Alignment/Map format and SAMtools. Bioinformatics 25, 2078–2079, doi:10.1093/bioinformatics/btp352 (2009).

65 Zhang, Y. et al. Model-based analysis of ChIP-Seq (MACS). Genome biology 9, R137, doi:10.1186/gb-2008-9-9-r137 (2008).

66 Carroll, T. S., Liang, Z., Salama, R., Stark, R. & de Santiago, I. Impact of artifact removal on ChIP quality metrics in ChIP-seq and ChIP-exo data. Frontiers in Genetics 5, doi:10.3389/fgene.2014.00075 (2014).

67 Stark, R. & Brown, G. DiffBind: differential binding analysis of ChIP-Seq peak data. http://bioconductor.org/packages/release/bioc/vignettes/DiffBind/inst/doc/DiffBind.pdf. (2011).

68 Love, M. I., Huber, W. & Anders, S. Moderated estimation of fold change and dispersion for RNA-seq data with DESeq2. Genome biology 15, 550, doi:10.1186/s13059-014-0550-8 (2014).

69 Quinlan, A. R. & Hall, I. M. BEDTools: a flexible suite of utilities for comparing genomic features. Bioinformatics 26, 841–842, doi:10.1093/bioinformatics/btq033 (2010).

70 Yu, G., Wang, L.-G. & He, Q.-Y. ChIPseeker: an R/Bioconductor package for ChIP peak annotation, comparison and visualization. Bioinformatics 31, 2382–2383, doi:10.1093/bioinformatics/btv145 (2015).

71 McLean, C. Y. et al. GREAT improves functional interpretation of cis-regulatory regions. Nature Biotechnology 28, 495–501, doi:10.1038/nbt.1630 (2010).

72 Bailey, T. L. et al. MEME Suite: tools for motif discovery and searching. Nucleic acids research 37, W202–W208, doi:10.1093/nar/gkp335 (2009).

