## Supplemental Information for "Chromatin alterations in the aging lung change progenitor cell activity"

Figure S1.

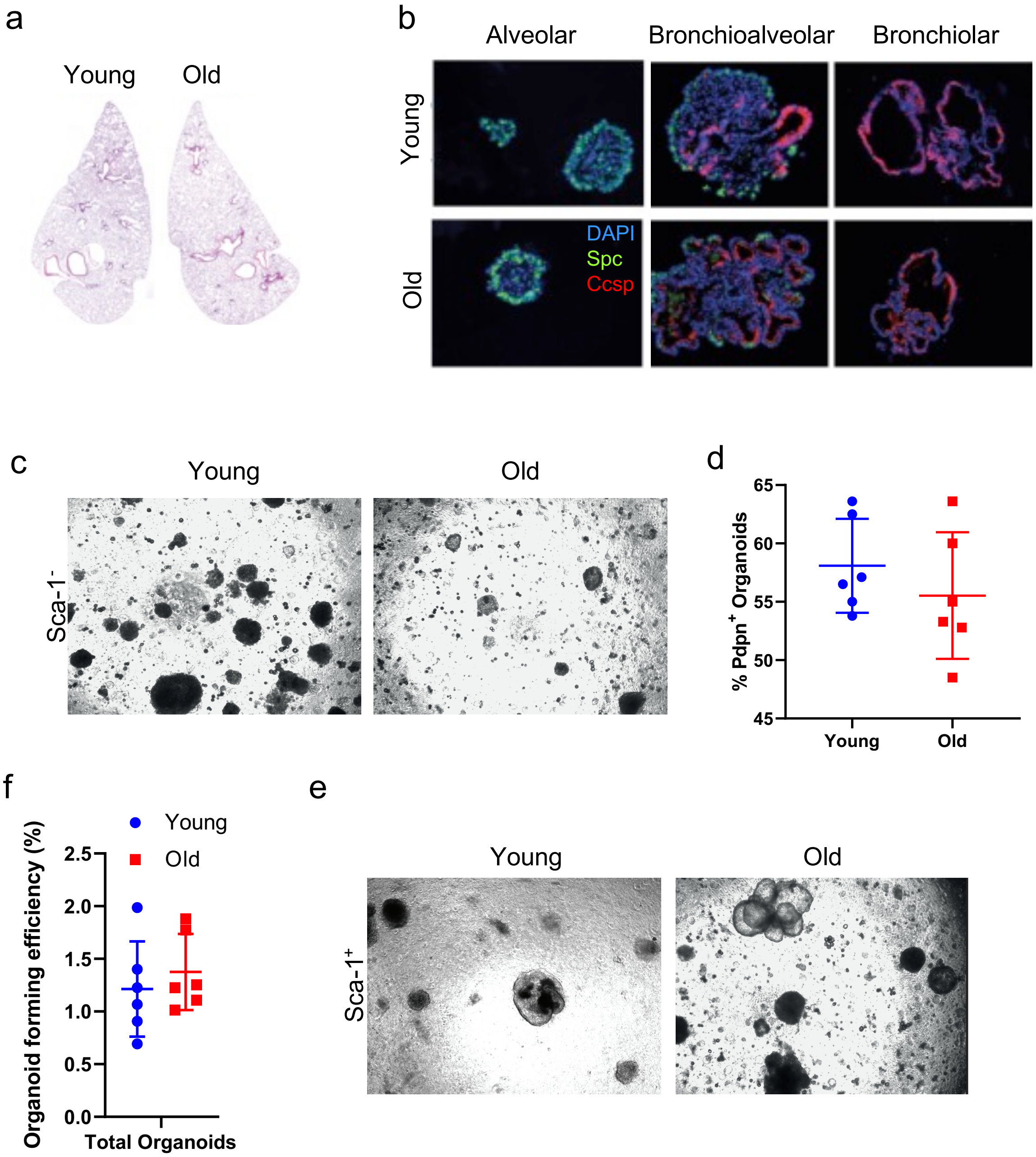

**Figure S1. (a)** Representative H&E stained images of whole lung lobes from young and old mice. **(b)** Representative images of major organoid types derived from Sca-1$^+$ lung progenitors from young and old mice, immunostained for the indicated proteins. **(c)** Representative brightfield images of organoids derived from Sca-1$^-$ lung progenitors from young and old mice. **(d)** Quantification of fraction of organoids derived from Sca-1$^-$ progenitors from young and old mice with Pdpn$^+$ cells. **(e)** Quantification of the organoid forming efficiency of Sca-1$^+$ progenitors isolated from young and old mice. **(f)** Representative brightfield images of organoids derived from Sca-1$^+$ lung progenitors from young and old mice.

Figure S2.

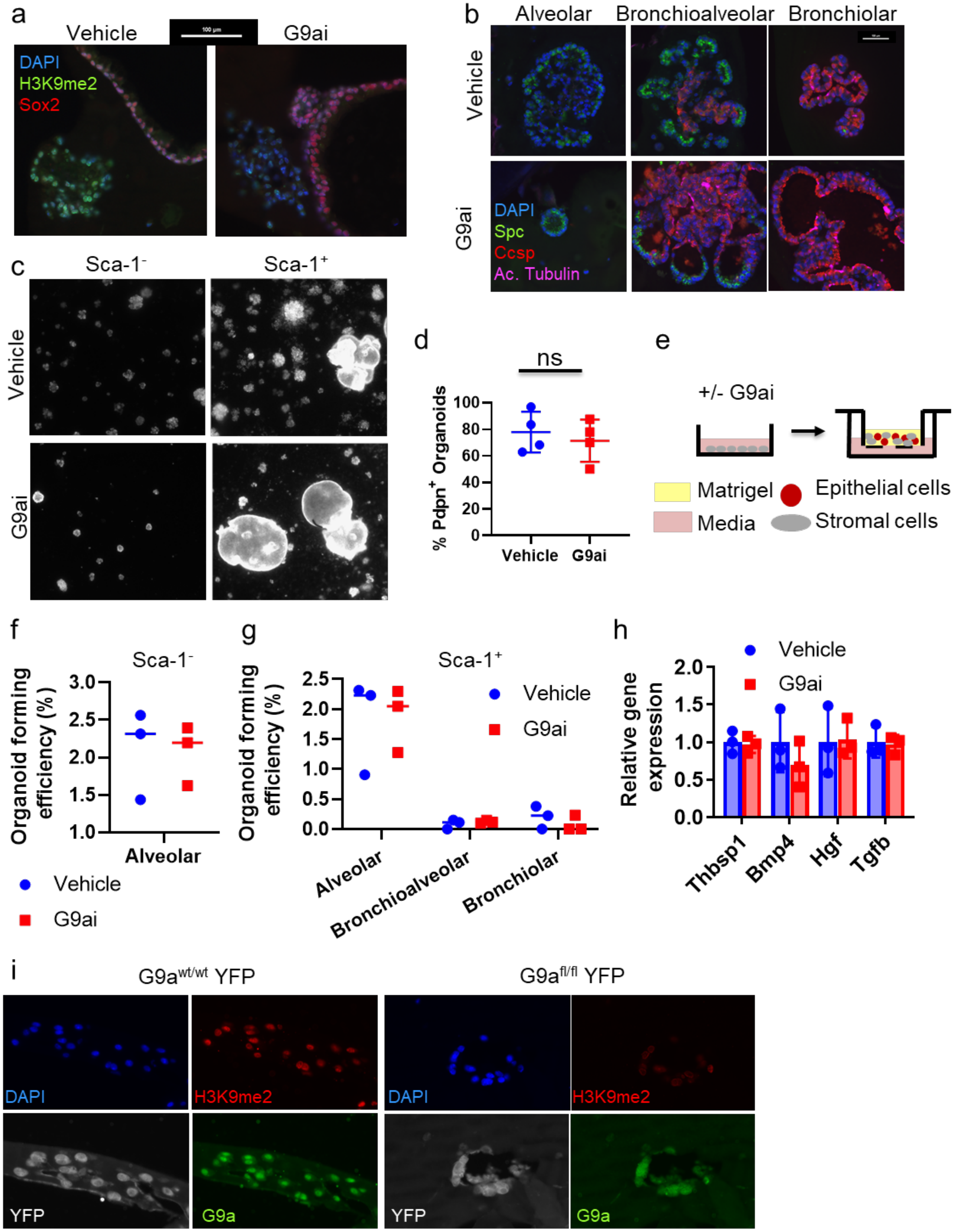

a
Vehicle
G9ai
DAPI
H3K9me2
Sox2
b
Alveolar
Bronchioalveolar
Bronchiolar
Vehicle
G9ai
DAPI
Spc
Ccsp
Ac. Tubulin
c
Sca-1⁻
Sca-1⁺
Vehicle
G9ai
d
ns
% Pdpn⁺ Organoids
100
80
60
40
20
0
Vehicle
G9ai
e
+/- G9ai
Matrigel
Epithelial cells
Media
Stromal cells
f
Sca-1⁻
Organoid forming efficiency (%)
3.0
2.5
2.0
1.5
1.0
Alveolar
Vehicle
G9ai
g
Sca-1⁺
Organoid forming efficiency (%)
2.5
2.0
1.5
1.0
0.5
0.0
Vehicle
G9ai
Alveolar
Bronchioalveolar
Bronchiolar
h
Vehicle
G9ai
Relative gene expression
2.0
1.5
1.0
0.5
0.0
Thbsp1
Bmp4
Hgf
Tgfb
i
G9a wt/wt YFP
G9a fl/fl YFP
DAPI
H3K9me2
YFP
G9a

**Figure S2 (a)** Representative images of day 14 organoids +/- G9ai, immunostained for the indicated proteins. Scale bar =100 μM **(b)** Representative images of major organoid types derived from Sca-1$^{+}$ lung progenitors +/- G9ai. **(c)** Representative fluorescent Images of day 14 organoid cultures +/- G9ai. Scale bar =100 μM. Scale bar =100 μM . **(d)** Quantification of Pdpn+ organoid % relative to all alveolar organoids. Ns= not statistically significant, T-test. **(e)** Schematic of stromal cell in vitro G9ai experiment. **(f & g)** Generation efficiencies of major organoid types from Sca-1$^{-}$ and Sca-1$^{+}$ progenitors co-cultured stromal cells pre-treated +/- G9ai. ns=p>0.05, T -test. **(h)** Relative expression levels of indicated genes normalized to GAPDH in lung stromal cells +/- G9ai. **(i)** Representative images of organoids from Adeno-Cre infected Sca-1$^{-}$ progenitors from G9a$^{wt/wt}$ and G9a$^{fl/fl}$ mice, immunostained for the indicated proteins.

Figure S3.

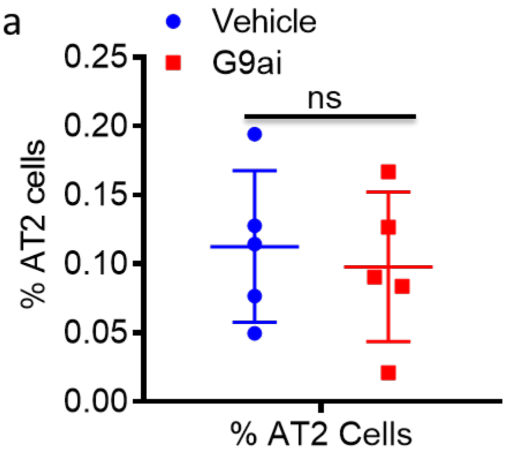


**Figure S3 (a)** Quantification of % AT2 over total cells per field in bleomycin damaged control and G9ai lungs. ns=p>0.05, T test.

Figure S4.

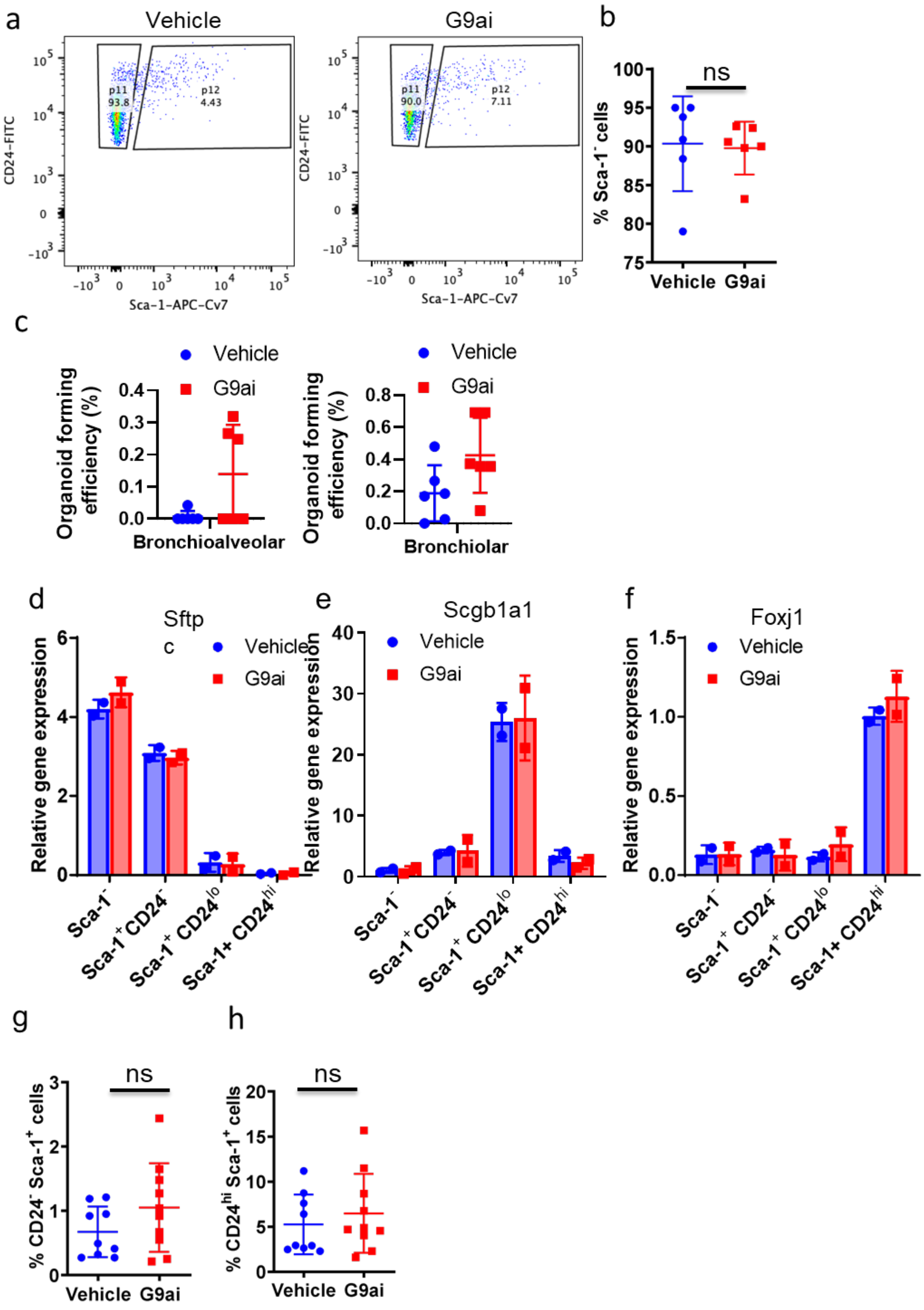

a
Vehicle
G9ai
CD24-FITC
Sca-1-APC-Cv7
p11 93.8
p12 4.43
p11 90.0
p12 7.11
b
ns
% Sca-1- cells
Vehicle G9ai
c
Vehicle
G9ai
Organoid forming efficiency (%)
Bronchioalveolar
Bronchiolar
d
Sftp c
e
Scgb1a1
f
Foxj1
Relative gene expression
Sca-1-
Sca-1+ CD24-
Sca-1+ CD24lo
Sca-1+ CD24hi
g
h
% CD24- Sca-1+ cells
% CD24hi Sca-1+ cells

**Figure S4 (a)** Representative FACs plots of G9ai and control isolated lungs, gated for single, live, CD31/CD45-, Epcam$^+$ cells. **(b)** Quantification of Sca-1$^-$ fraction of lung epithelial cells, gated for single, live CD31/CD45$^+$, Epcam$^+$ cells. ns=p>0.05, T test. test **(d-f)** GAPDH normalized gene expression levels of **(d)** Sftpc, **(e)** Scgb1a1 and **(f)** Foxj1 in Sca-1 and CD24 sorted lung epithelial cells from G9ai and control mice. **(g & h)** Quantification of the **(g)** Sca-1$^+$ CD24$^-$ and **(h)** Sca-1$^+$ CD24$^{hi}$ fractions of lung epithelial cells, gated for single, live CD31/CD45$^+$, Epcam$^+$ cells. ns=p>0.05, T-test. **(i)** Representative images of organoids derived from lung progenitors +/- in vivo G9ai following 14 days in culture, immunostained for the indicated proteins.

Figure S5.

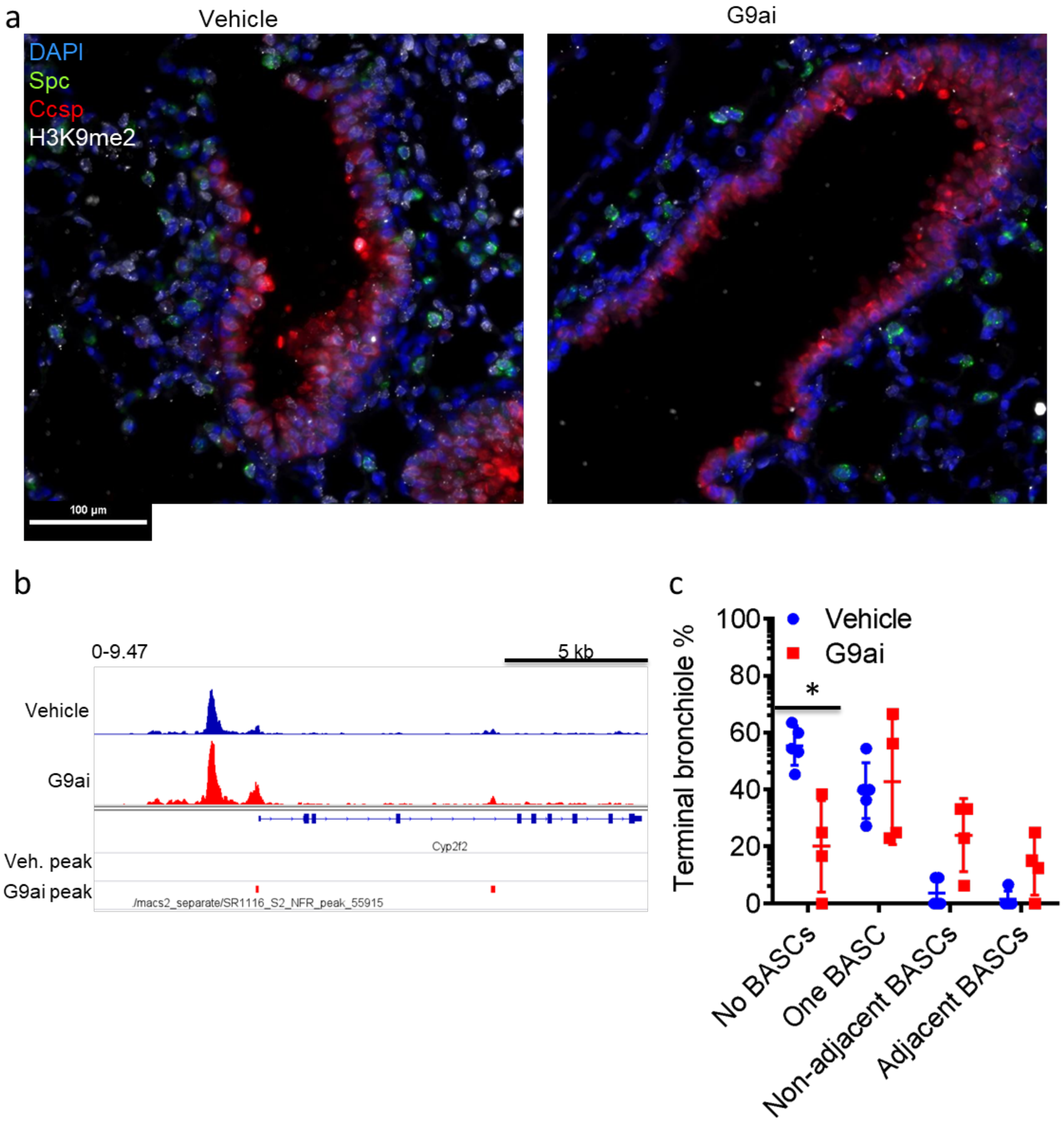

a
Vehicle
G9ai
DAPI
Spc
Ccsp
H3K9me2
100 µm
b
0-9.47
5 kb
Vehicle
G9ai
Cyp2f2
Veh. peak
G9ai peak
/macs2_separate/SR1116_S2_NFR_peak_55915
c
Vehicle
G9ai
Terminal bronchiole %
100
80
60
40
20
0
*
No BASCs
One BASC
Non-adjacent BASCs
Adjacent BASCs

**Figure S5 (a)** Representative images of day 3 terminal bronchioles in uninjured G9ai and control mice, immunostained for the indicated proteins. Scale bar = 100 μM **(b)** Quantification of % of terminal bronchioles with the indicated number and positions of BASCs in uninjured control and G9ai mice at day 3. *=$p<0.05$, Sidak's multiple comparison test. **(e)** Representative images of terminal bronchioles in G9ai and control mice. Arrows demark $Spc^+$ $Ccsp^+$ BASCs. Scale bar = 100 μM

Figure S6.

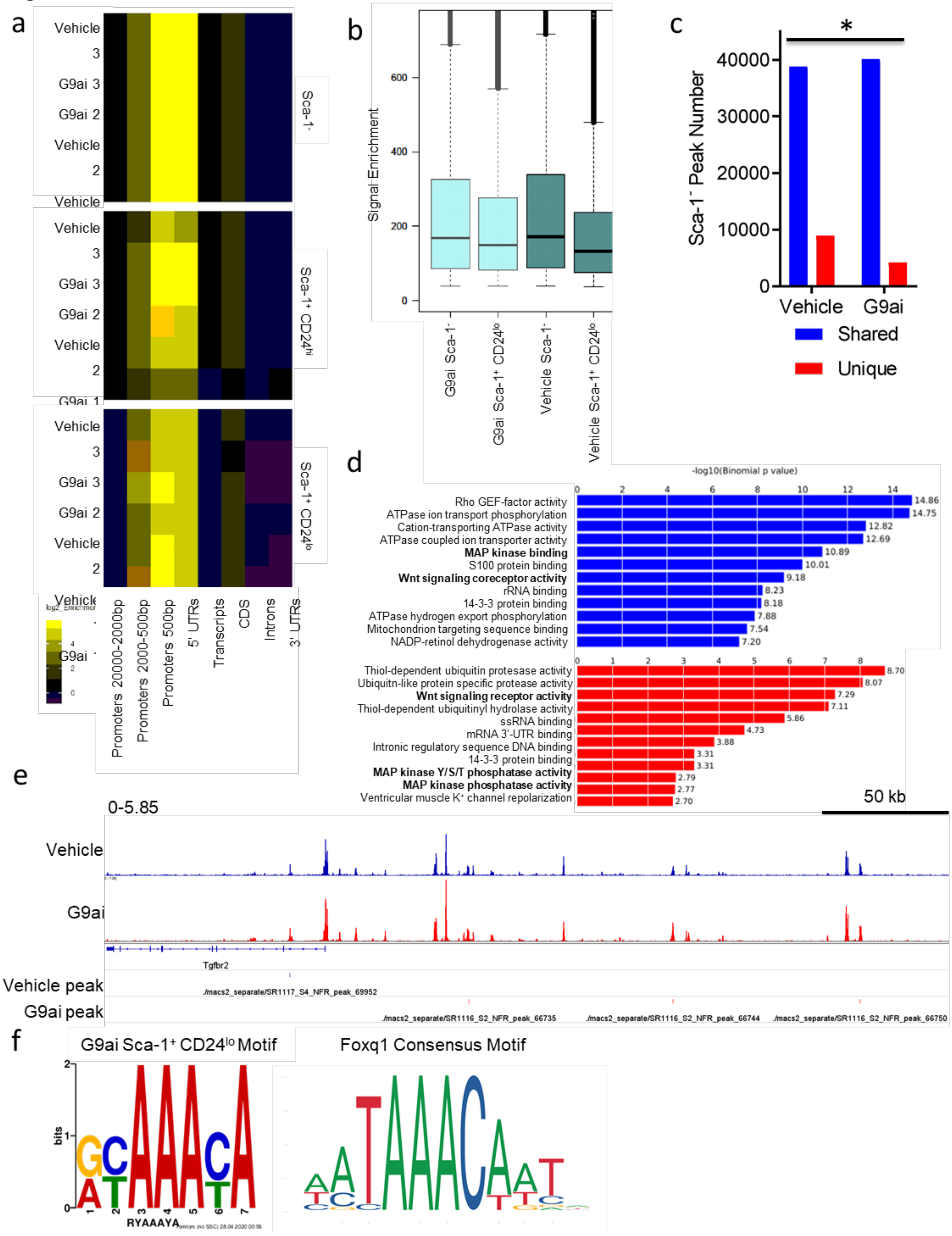
a
Vehicle 3
G9ai 3
G9ai 2
Vehicle 2
Vehicle
G9ai 1
Sca-1-
Sca-1+ CD24hi
Sca-1+ CD24lo
log2_Enrichment
Promoters 20000-2000bp
Promoters 2000-500bp
Promoters 500bp
5' UTRs
Transcripts
CDS
Introns
3' UTRs
b
Signal Enrichment
600
400
200
0
G9ai Sca-1-
G9ai Sca-1+ CD24lo
Vehicle Sca-1-
Vehicle Sca-1+ CD24lo
c
*
Sca-1- Peak Number
40000
30000
20000
10000
0
Vehicle
G9ai
Shared
Unique
d
-log10(Binomial p value)
Rho GEF-factor activity 14.86
ATPase ion transport phosphorylation 14.75
Cation-transporting ATPase activity 12.82
ATPase coupled ion transporter activity 12.69
MAP kinase binding 10.89
S100 protein binding 10.01
Wnt signaling coreceptor activity 9.18
rRNA binding 8.23
14-3-3 protein binding 8.18
ATPase hydrogen export phosphorylation 7.88
Mitochondrion targeting sequence binding 7.54
NADP-retinol dehydrogenase activity 7.20
Thiol-dependent ubiquitin protesase activity 8.70
Ubiquitn-like protein specific protease activity 8.07
Wnt signaling receptor activity 7.29
Thiol-dependent ubiquitinyl hydrolase activity 7.11
ssRNA binding 5.86
mRNA 3'-UTR binding 4.73
Intronic regulatory sequence DNA binding 3.88
14-3-3 protein binding 3.31
MAP kinase Y/S/T phosphatase activity 3.31
MAP kinase phosphatase activity 2.79
Ventricular muscle K+ channel repolarization 2.77
2.70
e
0-5.85
50 kb
Vehicle
G9ai
Tgfbr2
Vehicle peak
G9ai peak
./macs2_separate/SR1117_S4_NFR_peak_69952
./macs2_separate/SR1116_S2_NFR_peak_66735
./macs2_separate/SR1116_S2_NFR_peak_66744
./macs2_separate/SR1116_S2_NFR_peak_66750
f
G9ai Sca-1+ CD24lo Motif
Foxq1 Consensus Motif
bits
RYAAAYA

**Figure S6. (a)** Heatmap of ATAC reads categorized by distance to TSS from vehicle and G9ai Sca-1$^{--}$, Sca-1$^{+}$ CD24$^{lo}$ and Sca-1$^{+}$ CD24$^{hi}$ cells. **(b)** Boxplots of whole genome open chromatin signal from vehicle and G9ai Sca-1$^{--}$, Sca-1$^{+}$ CD24$^{lo}$ and Sca-1$^{+}$ CD24$^{hi}$ cells. **(c)** Quantification of common and unique ATAC-seq chromatin peaks in vehicle and G9ai Sca-1$^{-}$ lung epithelial cells, $p<0.05$, $\chi^2$ test. **(d)** Table of significantly enriched GO Molecular Function terms associated with unique chromatin peaks from vehicle (blue) and G9ai (red) Sca-1$^{-}$ cells. **(e)** Representative chromatin tracks of vehicle and G9ai Sca-1$^{+}$ CD24$^{lo}$ cells, with position of unique peaks from each cohort indicated. **(f)** Motif comparison between G9ai Sca-1$^{+}$ CD24$^{lo}$ unique peak enriched motif and Foxq1 consensus.

Figure S7.

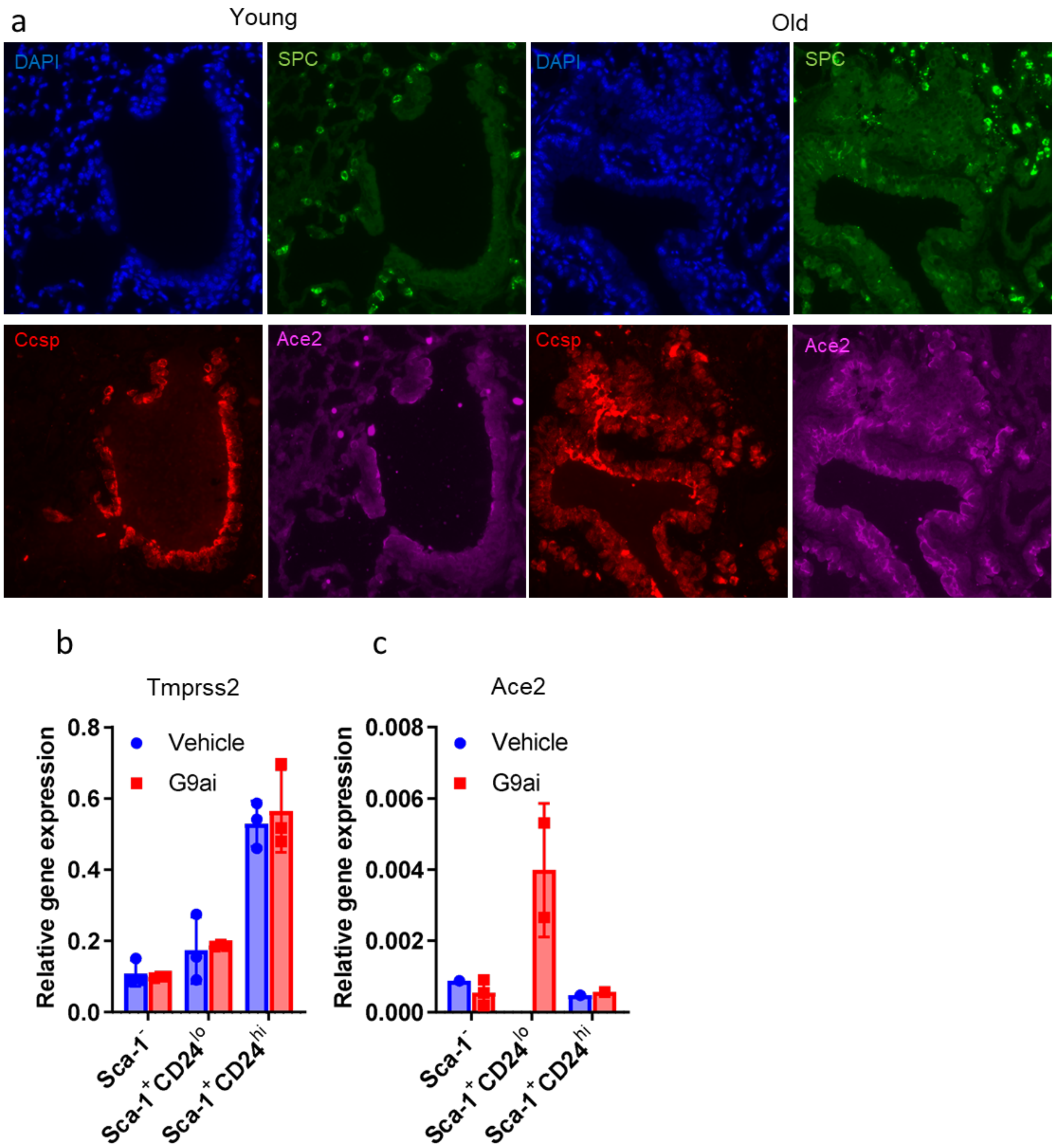


**Figure S7 (a)** Representative images of young and old lungs, immunostained for the indicated proteins **(b-c)** GAPDH normalized gene expression levels of **(a)** Tmprss2 and **(b)** Ace2 in Sca-1 and CD24 sorted lung epithelial cells from G9ai and control mice.

**Table S1.**

| Enriched G9ai Peak | Enriched Vehicle Peak | G9ai & Vehicle peaks | No Enriched peak |
|---|---|---|---|
| Scgb1a1 | Chad | Bpifb1 | Fmo2 |
| Scgb3a2 | Gdpd2 | Cldn10 | Pglyrp1 |
| Muc5b | 5330417C22Rik | Gabrp | Emp1 |
| Pigr | Selenbp1 | Scnn1b | Ascc2 |
| Ltf | Egr1 | Kitl | Gm26917 |
| Plpp3 | Lypd2 | Por | Fam46c |
| Fmo3 | Mgst1 | Epas1 | Hspa1b |
| Aldh3a1 | Adrb2 | Ifrd1 | Bpifa1 |
| Lrrc26 | | Klf4 | 8430408G22Rik |
| Tff2 | | Malat1 | Retnla |
| Adh7 | | Mecom | Hp |
| Kcnk2 | | Runx1 | Hspa1a |
| Synpo | | | Gsta3 |
| Samd4 | | | Foxq1 |
| Reg3g | | | Cckar |
| Gmnn | | | Reep6 |
| Cp | | | C3 |
| Scgb3a1 | | | AU021092 |
| Cyp2f2 | | | Idh1 |
| Cldn4 | | | Hspb1 |
| Pon1 | | | Rasd1 |

| | |
|---|---|
| **Slc16a11** | **Junb** |
| **Btg2** | **Aldh1a7** |
| **Slc4a5** | **Vars** |
| **Trf** | **Gstm2** |
| **Pam** | **Gm42418** |
| **Jun** | **Cyr61** |
| **Shh** | **Clu** |
| **Trim3** | **Ptgs2** |
| **Neat1** | **Wfdc2** |
| **Aox3** | **Aldh1a1** |
| **Timp3** | **Ier2** |
| **Osgin1** | **Qsox1** |
| **Scgb1c1** | **Gsto1** |
| **Hes1** | **2900060B14Rik** |
| **Fos** | **Muc1** |
| **Prdx6** | **Jund** |
| **Ahnak** | **Slc1a5** |
| **Krt18** | **Son** |
| **Fam13a** | **Rpsa** |
| **Aldh2** | **Rrbp1** |

**Table S1.** List of Club cell signature genes categorized by the presence of enriched ATAC-seq peaks in G9ai and Vehicle Sca-1 CD24$^{lo}$ bronchiolar progenitor cells.

### Table S2

| Enriched G9ai Peak | | | Enriched Vehicle Peak | G9ai & Vehicle peaks | No enriched peak | | | | |
|---|---|---|---|---|---|---|---|---|---|
| Sftpa1 | Slc38a2 | Fam102b | Scd1 | Sftpd | Lyz2 | Lrp2 | Rplp0 | Renbp | Npnt |
| Cxcl15 | Sptlc2 | Slc25a51 | Slc34a2 | H2-Aa | Sftpc | Glrx | Rps15 | Chd3 | Car2 |
| Sfta2 | Antxr1 | Slc43a2 | S100g | Ctsc | Lyz1 | Slco4c1 | Rps21 | Stard3nl | Smim1 |
| Tmsb4x | Rpl19 | Dennd5a | Cd74 | Fgfr2 | Chil1 | Eef2 | Mtch1 | Mthfd1 | Bmp4 |
| H2-Ab1 | Tns1 | Myof | Lcn2 | Eef1a1 | Fabp5 | Cnn2 | Rpl30 | Slc26a9 | Sec14l4 |
| Bex2 | Dbi | H2-D1 | Rnase4 | App | Sftpb | Rpl36a | H2-DMa | Pdzk1ip1 | Asah1 |
| Elovl1 | Rbms3 | Dram2 | Il33 | Rps14 | Napsa | Tmem30a | Adamts1 | Eif3e | Hnrnpa1 |
| Cd36 | Lcp1 | Sox4 | Tmem243 | Pi4k2b | Npc2 | Rpl14 | Rps11 | Bex1 | Pank3 |
| Apoc1 | Man1a | Pdlim2 | Aqp5 | Kcnj15 | Lamp3 | Use1 | Dpp4 | St6galnac4 | Gadd45a |
| Errfi1 | AA986860 | Eif3k | Rasl11a | Phldb2 | Lpcat1 | Rpl37 | Sgpp2 | Gpx1 | Wsb1 |
| H2-Eb1 | Rai14 | Hnmt | Adgrf5 | Ahcyl2 | Bex4 | Bcam | Rps29 | Vasn | Zfp36l2 |
| Oat | Aox3 | Rps6ka3 | Gem | Fosb | Hc | Exosc7 | Fau | Cyp4v3 | Fam126b |
| Irx1 | Esam | Hmgcs1 | Tpt1 | Secisbp2l | Dram1 | Nrn1 | Rps25 | Mtcl1 | Ece1 |
| Acsl4 | Ptgfrn | Nedd4l | Rpl7 | Ctsh | Pla2g1b | Dusp1 | Vsig2 | Tubb5 | Slc31a1 |
| Nrp1 | Trf | Anxa6 | Gja1 | Car8 | Sepp1 | Ociad2 | Naca | Chp1 | Rpl36al |
| Acot7 | Snhg18 | Csgalnact2 | Nkd1 | Sgms1 | Rgcc | Prps2 | Col6a1 | Ddx3x | 2700094K13Rik |
| Abca3 | Tfcp2l1 | Nr3c1 | Rps15a | Gclc | Ager | Ang | Slpi | Maob | Cyp51 |
| Zdhhc3 | Vamp8 | Arpc1b | Rbpjl | Ank3 | Rps19 | Rps20 | Rpl9 | Usp8 | Ppp1r10 |
| Stmn1 | Adcy7 | Polr2e | Spry1 | Ctnnb1 | Lrg1 | Sparc | Rps17 | Ddhd1 | Tpp1 |
| Cited2 | Jun | Lamp2 | S100a10 | Ccdc141 | Etv5 | Snx7 | Rgs10 | Tnfaip1 | Tmem50b |
| Matn4 | Zfp36 | Tuba1c | Ccz1 | Chsy1 | Fasn | Atp6v1c2 | Rpl39 | Gpt | Epha2 |
| Atp8a1 | Fos | Ddx6 | Anpep | Iqgap1 | Lgi3 | Eef1b2 | Tmem110 | Arg2 | Plin2 |
| Cd200 | Rpl11 | Rassf3 | Ptgs1 | Sdc1 | Tgoln1 | Rpl7a | Rps4x | Rps13 | Cpeb4 |
| Slc39a8 | Rpl36 | Stx12 | Bmp1 | Anxa3 | Apoe | Fdps | Pvrl1 | Atp6v1b2 | Adssl1 |
| Phlda1 | Eif3h | Srebf1 | Limd2 | Npc1 | Egfl6 | Cst8 | Rpl35a | Azin1 | Plekhb2 |

| | | | | | | | | | |
|---|---|---|---|---|---|---|---|---|---|
| Cd63 | Cat | Akap7 | Prrg3 | Ppp1r9a | Cldn18 | Id2 | Rplp2 | Csrp1 | Rps12-ps3 |
| Soat1 | Cadm1 | Scamp1 | Tmem159 | Itga9 | Scd2 | Rps6 | Rpl23 | Syngr1 | D230025D16Rik |
| Irx2 | Rmdn2 | Imp3 | Krt23 | Epas1 | Cox6a2 | Rpl27a | Rpl35 | Unc119 | 1110008P14Rik |
| Atp11a | Gprc5a | Ctsb | Ggcx | Me1 | Acly | Rpl34 | Junb | Urah | Eif3m |
| Igfbp7 | Slc16a1 | Hspb8 | Apc | Irf2bp2 | Muc1 | Rpl18 | Ier2 | Clk1 | Deptor |
| Rps23 | Scgb1c1 | Dap | Acaca | Mylk | Rpl21 | Rpl4 | Rpl31 | Usp33 | Gstt3 |
| Abhd2 | Arl4d | | Lamb3 | Kcne2 | Enpep | Idi1 | Rpsa | Lgals9 | Grb14 |
| Slc12a2 | Slc1a3 | | Dlc1 | Rps26 | Rps28 | Irs2 | mt-Nd3 | Hmox1 | Idh1 |
| Ramp1 | Adgrg1 | | Nipal1 | Gsap | Rpl10 | H2-DMb1 | Rpl6 | Marveld1 | Atf4 |
| Prnp | Wls | | Spry2 | Lrrc8c | Rps12 | Sat1 | mt-Co2 | Fam101b | Ngfrap1 |
| Cebpa | Rpl38 | | Gadd45b | Col4a1 | Wbp5 | Adam19 | Tceal8 | Epn3 | Ptgs2 |
| Tfrc | 9230117E06Rik | | Ly6c2 | Fads1 | Npw | Hk2 | Rps2 | Hexim1 | |
| Snx25 | Brd7 | | Igsf3 | Vegfa | Acoxl | Rpl22 | Sertad1 | Stbd1 | |
| Taok3 | Eif3f | | Trib1 | Fam178b | Mme | Gnb2l1 | Cyba | H1f0 | |
| Ppp1r14c | Pnpla2 | | Ehf | C6 | Rpl18a | Eef1g | Rpl24 | Gm11744 | |
| Rab27b | Rtkn | | Hacd3 | Tc2n | Cpm | Rps18 | Entpd1 | Scarb2 | |
| Degs1 | Tspan12 | | Nkx2-1 | Rerg | Adk | Rpl10a | Tmem42 | Pgm2 | |
| Lrrk2 | Abhd5 | | Ubr1 | Rora | Chia1 | Rpl26 | Serpinb6b | Serpinb9 | |
| Arf6 | Bcl2a1b | | Sdc4 | St3gal1 | H2afj | Ctsz | Rpl12 | Pik3ca | |
| Rps24 | Ttll7 | | Stat3 | Slc22a23 | Cd81 | Insig1 | Igfbp6 | Atp6v0a1 | |
| Rps3 | Prmt8 | | Traf1 | Pfkfb4 | Mlc1 | Rpl32 | Rprm | Acat2 | |
| Pmvk | Mbnl1 | | Cndp2 | Snx30 | Rpl23a | Hnrnph1 | Bcl2l11 | Pvrl2 | |
| Rps5 | Lmo4 | | Tmem97 | Mtss1 | Rpl3 | Kcnc3 | Sowahc | Nufip2 | |
| Tspan11 | Akr1b3 | | Arhgap44 | Foxo3 | Rps27a | Rpl17 | Cyb5b | Alad | |
| Nfkbiz | Fabp12 | | Snx4 | Klf4 | Icam1 | Chka | Epha7 | Rpl22l1 | |
| Gstt1 | Cd44 | | Ralgapa2 | Gadd45g | Bambi | H2-K1 | Snhg11 | Grn | |
| Acsl5 | Pabpc1 | | Abracl | Pim3 | Iah1 | Cers2 | Atp6v1a | Snhg12 | |
| Mal | Tspan8 | | Capg | P4ha1 | Zfos1 | Rpl23a-ps3 | Col4a2 | Yae1d1 | |

| | | | | | | | |
|---|---|---|---|---|---|---|---|
| **Socs2** | **Mob1b** | **Mtfr1l** | **Btg3** | **Mid1ip1** | **Rab9** | **Jund** | **Rap1gap** |
| **Alcam** | **Spr** | | **Zfp36l1** | **Rpl5** | **Rps27** | **Dnajb9** | **Gfpt1** |
| **Tmem164** | **Ube2i** | | **Hbegf** | **Ptprf** | **Rps16** | **H6pd** | **Hsd17b10** |
| **Rpl37a** | **Gata6** | | **Sort1** | **Rps3a1** | **Mbip** | **Cbx4** | **Clu** |
| **Abcd3** | **Spred1** | | **Tsc22d3** | **Scp2** | **Tinag** | **Tuba4a** | **Dag1** |
| **Nckap5** | **Mdfic** | | **Frmd6** | **Atp1b1** | **Zfp503** | **St6galnac2** | **Tnfsf9** |
| **Met** | **Pon3** | | **Fndc3b** | **Rps9** | **Tinagl1** | **Tob2** | **Mcfd2** |
| **Cox7a2l** | **Ccng2** | | **Hpcal1** | **Rps7** | **Rpl8** | **Fam173a** | **Fzd5** |
| **Nucb2** | **Camk2n1** | | **Fut8** | **Ppic** | **Rab27a** | **Cd302** | **Cxcl16** |
| **Mcl1** | **Lpin2** | | **Mapkapk2** | **Slco2a1** | **Tnfrsf12a** | **Marcksl1** | **Herpud1** |
| **Tmem163** | **Kif13a** | | **Nfix** | **Rpl13a** | **Rps10** | **Gltscr2** | **Nnat** |
| **Ldlr** | **Arhgef38** | | **Bhlhe40** | **Itih4** | **Rpl13** | **Paqr8** | **Brd2** |

**Table S2.** List of AT2 cell signature genes categorized by the presence of unique ATAC-seq peaks in G9ai and Vehicle Sca-1 CD24$^{lo}$ bronchiolar progenitor cells.
